# Functional connectivity during drawing after upper extremity peripheral nerve surgery: enhanced connectivity between motor and visuomotor-parietal regions

**DOI:** 10.64898/2026.04.28.721485

**Authors:** S Gassass, MD Wheelock, N Kapil, T Kim, David M. Brogan, Christopher J. Dy, Susan E. Mackinnon, BA Philip

**Author notes:** **Corresponding Author: Samah Gassass, MS, OTR/L**, Department of Occupational Therapy, Washington University School of Medicine in St. Louis, 4444 Forest Park Avenue, St. Louis, MO 63108, USA, Email (primary), Email (personal).

## Abstract

**Importance:** Recovery after upper extremity peripheral nerve injury (PNI) surgery depends on changes in cortical neural patterns that support sensorimotor control. Task-based functional connectivity (FC) can characterize these changes, yet few studies have explored FC during ecologically fine motor valid tasks after PNI.

**Objective:** To investigate task-based FC with the left primary motor cortex (M1) during right hand drawing in individuals following right hand PNI surgery.

**Participants:** Forty-four right-handed adults, including 12 patients post PNI surgery (n = 8 with nerve repair, n = 4 with nerve transfer) and 32 healthy controls.

**Methods:** All participants underwent fMRI while performing a RH visuomotor precision drawing task. Seed-based connectivity analysis was performed to characterize the pattern of FC between left M1 and all voxels in the brain. We hypothesized that left M1 FC would differ between patients and controls, between Repair and Transfer groups, and covary with time since surgery.

**Results:** Patients (vs. controls) showed greater FC between left M1 and right visual and premotor cortices. Nerve transfer (vs. repair) showed greater FC between left M1 and right inferior parietal areas. Time since surgery was not linearly related to FC, though exploratory analyses suggested a negative association between log-time and FC between left M1 and right inferior parietal lobule.

**Conclusion:** After PNI surgery, visuomotor precision drawing involved distinct and behaviorally relevant neural patterns, which varied by task demand and potentially by surgical group despite clinical heterogeneity. Inferior parietal cortex may be especially engaged in early months after surgery (i.e. log-time). To improve recovery of upper limb function after PNI, clinical recommendations include incorporating early function-specific dexterous training, tailoring rehabilitation across surgical and recovery stages, and using multidimensional assessments of hand function.

## Introduction

Peripheral nerve injury (PNI) in the upper extremity can result in substantial sensorimotor impairment (Isaacs, 2013; Taylor et al., 2009), but little is known about how cortical plasticity supports functional recovery after treatment of PNI. Initial PNI management often involves nerve repair (“Repair” for brevity) such as direct coaptation (Pereira et al., 2023), nerve grafting (Lee & Wolfe, 2000), release or decompression (Khan & Perera, 2020) when the affected nerve(s) remain viable (Kline, 1990). Alternatively, more complex reconstruction such as nerve transfer (“Transfer” for brevity) may be performed in cases of extensive damage, delayed presentation, or suboptimal recovery following repairs (Tung & Mackinnon, 2010). Nerve transfer involves rerouting a functionally intact donor nerve to reinnervate the target muscles (Moore, 2014). Although repair and transfer differ in surgical technique and the cortical adaptation demands they trigger (Navarro, 2009; Navarro et al., 2007), both ultimately depend on the brain’s capacity to reestablish effective corticospinal and peripheral motor connectivity to regain volitional motor control of the reinnervated muscles (Kaas, 2000; Valyear et al., 2019).

Neuroimaging studies have shown that cortical plasticity after repair and transfer can involve multiple, distinct forms of neural changes, including altered somatotopic representation (Chen et al., 2013; Shen, 2022; Yao et al., 2015), functional remapping (Beaulieu et al., 2006), and/or the recruitment of additional cortical regions (Amini et al., 2018; Nordmark et al., 2018). When the cortical changes are maladaptive (e.g., aberrant sensorimotor reorganization) (Cramer et al., 2011; Kaas, 2000; Meyers et al., 2019), patients may experience limitations in the functional use of reinnervated muscles (Li et al., 2015; Machado et al., 2010) due to loss of selective muscle activation (Lemon, 2008). Therefore, characterizing the neural patterns associated with sensorimotor recovery is essential for developing targeted rehabilitation and neuromodulation interventions.

Task-based functional connectivity (FC) using functional magnetic resonance imaging (fMRI) offers an approach to characterize neural patterns after sensorimotor recovery. However, very few studies have investigated FC in humans after PNI surgery (Gassass, Lipsey, et al., 2025). One study demonstrated that early after intercostal-to-musculocutaneous (ICN-MCN) transfer, FC increased between the denervated arm brain area and the diaphragm representation during simple elbow flexion, a pattern associated with recovery of motor function after reinnervation (F. P. S. Fischmeister et al., 2020). Other resting-state FC studies after nerve transfers showed distributed cortical reorganization that lacked clear associations with clinical outcomes (Bhat et al., 2017; Fraiman et al., 2016; Yang, 2024; Yu et al., 2017). The few studies of task-based FC after PNI relied on constrained, single movement paradigms (e.g. Beisteiner et al., 2011; Florian Ph S Fischmeister et al., 2020; Sokki et al., 2012). Few if any studies have evaluated FC during ecologically valid dexterous movement, despite task-based FC’s high behavioral relevance, context specificity, and fewer confounds compared to resting-state FC (Vanderwal et al., 2019; Zhao et al., 2023).

The primary aim of this study was to investigate task-based FC patterns in patients following PNI surgical repair and transfer in the right hand. We assessed FC between left primary motor cortex (M1) and the rest of the brain during the right hand execution of a naturalistic visuomotor precision drawing task (PDT). We tested three hypotheses: (1) FC differs between PNI patients and healthy controls; (2) FC differs between Transfer and Repair groups; and (3) FC varies as a function of time since surgery in all PNI patients.

In secondary analyses, we compared each patient group with controls (Repair > Controls; Transfer > Controls) and examined whether individual differences in hand use and participation in precision-hand activities were associated with M1-related FC in all participants and separately within the patient group.

## Methods

### Study Sample and Recruitment

This cross-sectional study included 44 right-handed adults (29 females, 15 males, median age 40 [21–58]), including 4 patients with nerve transfer surgery (“Transfer”), 8 patients with nerve repair surgery (“Repair”), and 32 healthy controls (“Controls”). Demographic details are shown in **Table 1**. Between-group comparisons for categorical variables were performed using chi-square (χ^2^) tests, or Fisher’s exact tests when expected cell counts were < 5. Post-hoc pairwise comparisons were performed using Fisher’s exact tests with Holm correction. Injury characteristics and surgical details are provided in **Supplementary Tables 1-2**. Six additional participants were excluded from analysis, either due to impairment that made them unable to complete the task with the affected right hand (n = 3), or poor task compliance/excess motion (n = 3; for details see Kapil et al., 2025); these participants are not included in the final sample (N = 44).

**Table 1:** Demographics of Study Participants.

TASK CONNECTIVITY AFTER NERVE SURGERY
| Variable | Total<br>(n = 44) | Healthy Controls<br>(n = 32) | Repair<br>(n = 8) | Transfer<br>(n = 4) | Between-groups *p* |
| --- | --- | --- | --- | --- | --- |
| Age, *median* [range] | 40 [21-82] | 36 [21-82] | 52.5 [34-67] | 53 [33-75] | 0.23 |
| Gender |  |  |  |  | 0.01* |
| Male | 15 (34.1) | 8 (25) | 3 (37.5) | 4 (100) |  |
| Female | 29 (65.9) | 24 (75) | 5 (62.5) | 0 |  |
| Race** |  |  |  |  | 0.84 |
| Asian | 3 (6.8) | 3 (9.4) | 0 | 0 |  |
| Black or African American | 6 (13.6) | 5 (15.6) | 0 | 1 (25) |  |
| Hispanic or Latino | 1 (2.3) | 1 (3.1) | 0 | 0 |  |
| White | 32 (72.7) | 21 (65.6) | 8 (100) | 3 (75) |  |
| Multiple | 1 (2.3) | 1 (3.1) | 0 | 0 |  |
| Prefer not to say | 1 (2.3) | 1 (3.1) | 0 | 0 |  |
| Education Level |  |  |  |  | 0.57 |
| Post-college degree | 12 (27.3) | 9 (28.1) | 2 (25) | 1 (25) |  |
| College graduate | 20 (45.5) | 15 (46.9) | 2 (25) | 3 (75) |  |
| Some college | 6 (13.6) | 3 (9.4) | 3 (37.5) | 0 |  |
| High school or equivalent | 2 (4.5) | 2 (6.2) | 0 | 0 |  |
| Other | 2 (4.5) | 1 (3.1) | 1 (12.5) | 0 |  |
| No response | 2 (4.5) | 2 (6.2) | 0 | 0 |  |
| Marital Status |  |  |  |  | 0.43 |
| Divorced | 1 (2.3) | 1 (3.1) | 0 | 0 |  |
| Married | 28 (63.6) | 17 (53.1) | 7 (87.5) | 4 (100) |  |
| Significant Other | 4 (9.1) | 3 (9.4) | 1 (12.5) | 0 |  |
| Single | 10 (22.7) | 10 (31.2) | 0 | 0 |  |
| Widowed | 1 (2.3) | 1 (3.1) | 0 | 0 |  |
| Employment Status |  |  |  |  | 0.08 |
| Employed full-time | 29 (65.9) | 23 (71.9) | 4 (50) | 2 (50) |  |
| Employed part-time | 6 (13.6) | 4 (12.5) | 2 (25) | 0 |  |
| Retired | 6 (13.6) | 4 (12.1) | 0 | 2 (50) |  |
| Unemployed | 3 (6.8) | 1 (3.1) | 2 (25) | 0 |  |
| Note. Unless otherwise specified, all values “n (%)” |  |  |  |  |  |

The current participants were selected chronologically from participants in Clinical Trial NCT05207878, who were recruited from nerve surgeon referral, medical records, and community outreach email lists. Inclusion criteria for the current study included: right-handedness by self-report and Edinburgh Handedness Inventory (EHI) (Oldfield, 1971) score ≥ +40. Additional inclusion criteria for patients only: chronic (≥ 6 months) upper extremity PNI (localized cause, not distributed pathology), self-reported difficulty writing (Disabilities of the Arm, Shoulder and Hand (DASH) survey) (Beaton et al., 2001), right motor performance ≥1 SD below age-matched norms on the Box and Blocks test (Mathiowetz et al., 1985), and history of surgery (repair or transfer) for their PNI.

Exclusion criteria included: chronic pain unrelated to the PNI; right hand motor diagnoses not related to the PNI; and amputation of the thumb, index, or middle fingers. Additional exclusion criteria for Controls: any motor diagnosis affecting either hand in the past two years. All experimental procedures were approved by the research ethics committees at Washington University in Saint Louis. All participants with usable data were included in the analysis, except as specified above.

### fMRI Data Collection

Scans were performed on a Siemens PRISMA 3T MRI scanner. High-resolution T1-weighted structural images were acquired, using the 3D MP-RAGE pulse sequence: TR = 4500 ms, TE = 3.16 ms, TI = 1000 ms, flip angle = 8.0°, 256 x 256 voxel matrix, FOV = 256 mm, 176 contiguous axial slices, resolution: 1.0 x 1.0 x 1.0 mm. A T2-weighted image was also acquired at: TR = 3000 ms, TE = 409 ms, 256 x 256 voxel matrix, FOV = 256 mm, 176 contiguous axial slices, resolution: 1.0 x 1.0 x 1.0 mm. Spin echo field maps were also collected before the functional runs. Blood oxygen level–dependent (BOLD) EPIs for fMRI were collected using a T2*-weighted gradient echo sequence, a standard 64-channel birdcage radio-frequency coil, and the following parameters: TR = 662 ms, TE = 30 ms, flip angle = 52°, 72 x 72 voxel matrix, FOV = 216 mm, 60 contiguous axial slices acquired in interleaved order, resolution: 3.0 x 3.0 x 3.0 mm, bandwidth = 2670 Hz/pixel, multi-band acceleration = 6x. Siemens auto-align was run at the start of each session.

### Precision Drawing Task

Participants completed six BOLD fMRI scans of a precision drawing task (PDT) using the STEGA app (Philip & Frey, 2014; Philip & Frey, 2016; Philip et al., 2023), alternating between right hand scans and left hand scans. They traced hollow shapes as accurately and quickly as possible, with real-time video feedback via an MRI-compatible tablet using bluescreen technology (Karimpoor et al., 2015 and **Supplementary Figure 1**). Pen position was recorded at 60 Hz. Each scan followed a block design: 10 cycles of 15.2 seconds drawing (45 shapes) alternated with 15.2 seconds rest, plus initial and final rest, totaling 5:23 minutes (488 volumes: 230 drawing, 258 rest).

### Measures of Upper Limb Function and Use

Upper limb function was characterized via 5 measures: four self-report standardized surveys and one movement-based task. All measures were completed for all participants unless otherwise noted.

#### Edinburgh Handedness Inventory

Self-reported hand preference was measured using the EHI (Oldfield, 1971). All participants completed the EHI to assess “current” hand preference at the time of the study (EHI_current_). For patients, this measure reflects hand preference after injury and surgery. Patients were also asked to complete the EHI retrospectively to rate their pre-injury hand preference (EHI_pre-injury_).

To characterize changes in self-reported hand preference associated with post-surgery, the outcome measure *“hand preference shift”* was calculated for each patient as the difference between current and pre-injury scores (ΔEHI = EHI_current_ − EHI_pre-injury_). Negative ΔEHI values indicate a shift toward the unaffected LH, positive values indicate a shift toward the affected right hand, and a ΔEHI value of zero indicates no change in EHI score relative to the pre-injury assessment.

#### Disabilities of the Arm, Shoulder and Hand (DASH)

Self-reported upper-limb disability was measured in all participants using the 30-item Disabilities of the Arm, Shoulder and Hand (DASH) (Hudak et al., 1996). For the Controls, three items (22, 23, and 30) were excluded as these items are not applicable to individuals without upper limb injury. The outcome measure *“DASH_Ability”* was calculated by multiplying the total DASH disability score by −1 (so that higher scores = higher reported ability).

#### Motor Activity Log – Amount Scale

Self-reported everyday use of the affected upper limb was measured in patients only (n = 12) using the Motor Activity Log (MAL) 30-item structured interview (Uswatte et al., 2006). Data were used from the MAL’s Amount Scale, which is quantified using a 0−5 scale where higher scores reflect more functional use. The outcome measure “*MAL_Amount*” was calculated for each participant as the average Amount Scale score across all items. Additionally, participants were able to provide optional comments regarding the affected hand use.

#### Activity Card Sort – Precision Hand Activity

Self-reported participation in precision-hand activities was derived from a subset of the Activity Card Sort version 2 (ACS-2) (Anthony et al., 2024; Baum & Edwards, 2008) based on precision grip classification criteria (Cutkosky, 1989; Napier, 1956) and prior work (Anthony et al., 2024; Gassass, Zhou, et al., 2025). The precision hand activities subset “ACS_Hand” was identified by three raters who reviewed the full 97-item ACS-2 and identified 26 precision-hand activities (e.g., writing, hand crafts); see **Supplementary Table 3** for the list of precision hand activities. The outcome measure *“Retention”* was calculated from the ACS_Hand using the standard “percentage of retained” formula, in which higher scores = more activities retained (Katz et al., 2003):

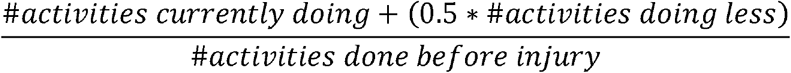

For Controls, the ACS questions were reworded to describe participation “in the past” instead of “before injury.”

#### Block Building Task (BBT) – Goal-Directed Right Hand Choices

To provide a quantitative assessment of right–left hand choices in an unconstrained goal-directed context, participants performed the Block Building Task (BBT) outside the scanner (Kim et al., 2025). Participants constructed LEGO models using both hands without instruction on which hand to use. Each participant completed two runs, and each run involved a table of 40 bricks without replacement. The outcome measure right hand use fraction *“RH_Use”* was calculated as the proportion of right grasps (injured hand grasps, in patients) out of total valid grasps, excluding invalid trials (e.g., scooping or dropping, representing 8% of the data). The “goal-directed” nature of the BBT reflects that the reach-to-grasp actions are not instructed but chosen by the participant in service of a goal (building a LEGO model).

RH use has a different meaning for different groups. In patients only, greater RH use would indicate functional recovery after right hand injury. In healthy adults, normative data indicate an average RH use of 0.63 (Kim et al., 2024; Stone & Gonzalez, 2015).

### Data Analysis

Descriptive statistical analyses were conducted separately for each measure of hand function: EHI_pre-injury_, EHI_current_, ΔEHI, DASH_Ability, MAL_Amount, Retention, and RH_Use. Descriptive statistics were reported as medians (ranges) and non-parametric tests were used to test group differences due to small sample sizes and/or non-normal distributions. On a post-hoc basis, correlations were assessed between months since surgery and a subset of the function variables (RH_Use, Retention, MAL_Amount, EHI_current_, ΔEHI), using Spearman’s ρ. Correction for multiple comparisons (15 correlations) was performed using Benjamini-Hochberg false discovery rate correction, with significance set at FDR-corrected α = 0.05, implemented in MATLAB 24.2.0 (Groppe, 2015).

For comparisons between patient groups (Repair vs Transfer; variables EHI and MAL_Amount), qualitative descriptions were used due to the small size of the Transfer group (n = 4); statistical analyses were used secondarily to quantify the observed effect, using Fisher’s exact test, Wilcoxon rank-sum tests (between-groups) or signed-rank tests (group vs. fixed value). For comparisons involving all three groups (Controls, Repair, Transfer; variables DASH_Ability, Retention, RH_Use), between-group differences were analyzed using Kruskal– Wallis tests or Fisher’s exact test for cells with counts <5. Significant omnibus effects were followed by pairwise group comparisons using Dunn’s or Fisher’s post-hoc tests with Holm correction. All tests were two-tailed α = 0.05 unless otherwise specified. Statistical analyses were conducted using R version 4.5.2 (R Foundation, Vienna, Austria).

### fMRI Analysis

#### fMRI Preprocessing

Functional and anatomical data were preprocessed using *fMRIPrep* 24.1.1 (O. Esteban et al., 2019; Esteban et al., 2018), which is based on *Nipype* 1.8.6 (Gorgolewski et al., 2011). Anatomical preprocessing included skull stripping, tissue segmentation, surface reconstruction (FreeSurfer 7.3.2), and normalization to MNI152NLin6Asym space at 2mm^3^ resolution using nonlinear registration (Team, 2020) with antsRegistration (ANTs 2.5.3) (Avants et al., 2011). Functional preprocessing included motion correction (FSL’s MCFLIRT) (Jenkinson et al., 2002), susceptibility distortion correction, coregistration to anatomical space (bbregister), and calculation of confounds including framewise displacement (FD) and three region-wise global signals (cerebrospinal fluid, white matter, and whole-brain). Additionally, physiological regressors were extracted to allow for component-based noise correction (CompCor) (Behzadi et al., 2007), with principal components estimated for temporal (tCompCor) and anatomical (aCompCor). The head-motion estimates and global signals were expanded with the inclusion of temporal derivatives and quadratic terms for each (Satterthwaite et al., 2013). Additional nuisance time series were calculated by means of principal components analysis of the signal found within a thin band (crown) of voxels around the edge of the brain (Patriat et al., 2017). All transformations were applied in a single resampling step using cubic B-spline interpolation. For full details on fMRIPrep preprocessing, see **Supplementary Text**.

Following fMRIPrep preprocessing, functional data underwent spatial smoothing with a 6 mm FWHM Gaussian kernel in CONN release 22.v2407 (Nieto-Castanon & Whitfield-Gabrieli, 2021).

#### Functional Connectivity (FC) Analysis

A seed-based connectivity (SBC, i.e. seed-to-voxel) analysis was performed to characterize the spatial pattern of FC between left M1 (contralateral to the moving RH, which was the injured hand in patients) and all voxels of the brain.

Functional data were denoised via CONN’s standard pipeline. Denoising included the regression of potential confounding effects characterized by 5 CompCor noise components each for cerebrospinal fluid and white matter (Behzadi et al., 2007; Chai et al., 2012), 12 motion regressors comprising the 6 rigid-body translation/rotation parameters and their first-order temporal derivatives (Friston et al., 1996), and up to 48 outlier scans (Power et al., 2014). Additional nuisance terms included 3 run/task effects and 2 linear trends. After regression, BOLD time series underwent high-pass frequency filtering above 0.008 Hz (Hallquist et al., 2013). The effective degrees of freedom of the BOLD signal after denoising were estimated to range from 2407.2 to 2713 (average 2599.4) across all subjects (Nieto-Castanon, 2025).

First-level (within-subjects) SBC were conducted in CONN using a seed in the left M1 normative hand region, defined as a 5 mm radius sphere centered on Montreal Neurological Institute (MNI) coordinates [X = −38 Y = −24, Z = 54] (Smith & Frey, 2011).

FC strength was represented by Fisher-transformed bivariate correlation coefficients from a weighted general linear model (weighted-GLM) (Nieto-Castanon, 2020), which models condition-specific associations between the seed and voxel-wise BOLD time series. The Draw condition was modeled as a boxcar function convolved with SPM’s canonical hemodynamic response function. FC during the Draw condition was analyzed directly, rather than using a Draw > Rest contrast, because the Rest condition exhibited widespread, non-specific correlations that reduced the interpretability of differential connectivity effects. Group-level analyses were performed using a GLM model (Nieto-Castanon, 2020). First-level M1 SBC maps served as dependent variables. Voxel-level effects were evaluated using multivariate parametric statistics with random-effects across subjects and sample covariance estimation across the three runs. Cluster-level inferences were based on parametric statistics from Gaussian Random Field theory (Nieto-Castanon, 2020; Worsley et al., 1996). To test the hypotheses, the following *a priori* group-level analyses were performed: group effects for Patients > Controls (hypothesis 1), group effects for Transfer > Repair (hypothesis 2), and covariate effects of time since surgery in patients (hypothesis 3).

To report each cluster, MNI coordinates were reported for the peak voxel, and region labels were determined for: (1) the peak voxel, (2) any anatomical areas covered by ≥10% of the cluster volume, and (3) any anatomical areas ≥10% covered by the cluster. Region labels were defined as the highest-probability gray matter region, plus any other gray matter region(s) with probability >25%, using the Juelich histological atlas (Amunts et al., 2020; Eickhoff et al., 2007). For regions without Juelich labels, the Harvard–Oxford atlas (Fischl et al., 2002) or MNI152 cerebellar atlas was used instead. Each cluster then received a name to briefly summarize the regions included.

Results were thresholded using a cluster-size threshold of familywise corrected p-FDR < 0.05 (Chumbley et al., 2010) and a voxel-level threshold of p < 0.01 (determined on a post-hoc basis as described below). For post-hoc determination of optimal voxel-level statistical thresholds for our small sample, the connectivity maps were examined at multiple voxel-level thresholds (*p* < 0.001, *p* < 0.01, *p* < 0.05) to identify the most stringent threshold that would reveal spatial patterns of FC across hypotheses 1 and 2, without producing the brain-wide clusters that limit interpretation in the absence of voxel-level thresholding (Woo et al., 2014). This resulted in the threshold of *p* < 0.01 as mentioned above. The cluster-size threshold was not adjusted in this process.

Due to negative results of “time since surgery,” the time analysis was expanded on a post-hoc basis to examine two non-linear time effects. First, to increase sensitivity to early changes, log transformed months since surgery (log months) was used. Second, to increase sensitivity to curvilinear relationships, quadratic time (months^2^) was used.

#### Exploratory analyses: Patient Group Differences and Effects of Retention and Hand Use on M1 Connectivity

Two exploratory analyses were performed to clarify and extend the findings of our main analyses. As in our main analyses, the exploratory analyses evaluated SBC with left M1 during right hand drawing. First, to determine whether one patient group drove the Patient > Control differences in SBC with left M1, we conducted post-hoc exploratory SBC analyses of the contrasts Repair > Controls and Transfer > Controls. The overlap between Patient > Control clusters and (Repair/Transfer) > Control clusters were reported as the percentage of voxels in the smaller cluster that were also inside the larger cluster.

Second, to explore relationships between SBC and behavior outside fMRI we tested the effects of Retention and RH_Use in all participants, and separately within the patient group only.

## Results

### Measures of Hand Function and Use in Daily Life

Participants completed four self-report measures and one movement-based task to assess upper-limb function and use following peripheral nerve injury (PNI) surgery. Individual scores are reported in **Table 2** and analyzed in the following sections.

**Table 2.** Individual participant measures of hand function and use.

TASK CONNECTIVITY AFTER NERVE SURGERY
| ID | Group | Mo. Since Injury | Mo. Since Surgery | EH<sub>I</sub><sub>pre-injury</sub> | EH<sub>I</sub><sub>current</sub> | ΔEH<sub>I</sub> | DASH Ability | MAL Amount | Retention (%) | RH Use |
| --- | --- | --- | --- | --- | --- | --- | --- | --- | --- | --- |
| 1004 | Repair | 425 | 418 | 100 | 30 | -70 | 54.16 | 3.61 | 26.66 | 0.58 |
| 1007 | Repair | 174 | 6 | 65 | 30 | -35 | 86.66 | 4.92 | 58.33 | 0.58 |
| 1011 | Repair | 5 | 2 | 80 | 80 | 0 | 92.24 | 4.65 | 58.33 | 0.61 |
| 1026 | Repair | 50 | 26 | 80 | 10 | -70 | 48.33 | 4.06 | 84.62 | 0.41 |
| 1032 | Repair | 73 | 12 | 100 | 85 | -15 | 89.16 | 3.78 | 100 | 0.51 |
| 1036 | Repair | 112 | 23 | 100 | -15 | -115 | 67.5 | 2.42 | 66.66 | 0.17 |
| 1038 | Repair | 11 | 1 | 100 | 40 | -60 | 73.33 | 4.72 | 43.75 | 0.65 |
| 1051 | Repair | 3 | 3 | 85 | 10 | -75 | 79.16 | 2.28 | 81.25 | 0.05 |
| 1009 | Transfer | 10 | 2 | 95 | 95 | 0 | 89.16 | 5 | 100 | 0.75 |
| 1023 | Transfer | 158 | 158 | 100 | 100 | 0 | 77.5 | 4.5 | 72.72 | 0.40 |
| 1027 | Transfer | 38 | 30 | 65 | 90 | 25 | 69.16 | 3.4 | 28.57 | 0.76 |
| 1046 | Transfer | 14 | 3 | 75 | -15 | -90 | 75 | 3.92 | 100 | 0.5 |

Note. Mo. = months. EH<sub>I</sub> = Edinburgh Handedness Inventory; ΔEH<sub>I</sub> = current – pre-injury. DASH = Disabilities of the Arm, Shoulder and Hand. MAL = Motor Activity Log. Retention = Activity Card Sort precision hand activities retention. RH Use = Block Building Task frequency of right hand choices.

#### Hand preference shift is significantly high in repair patients

Controls demonstrated strong right-hand (RH) preference at the time of the study (median EHI 97.5 [50–100]) (**Figure 1A**).

**Figure 1.**
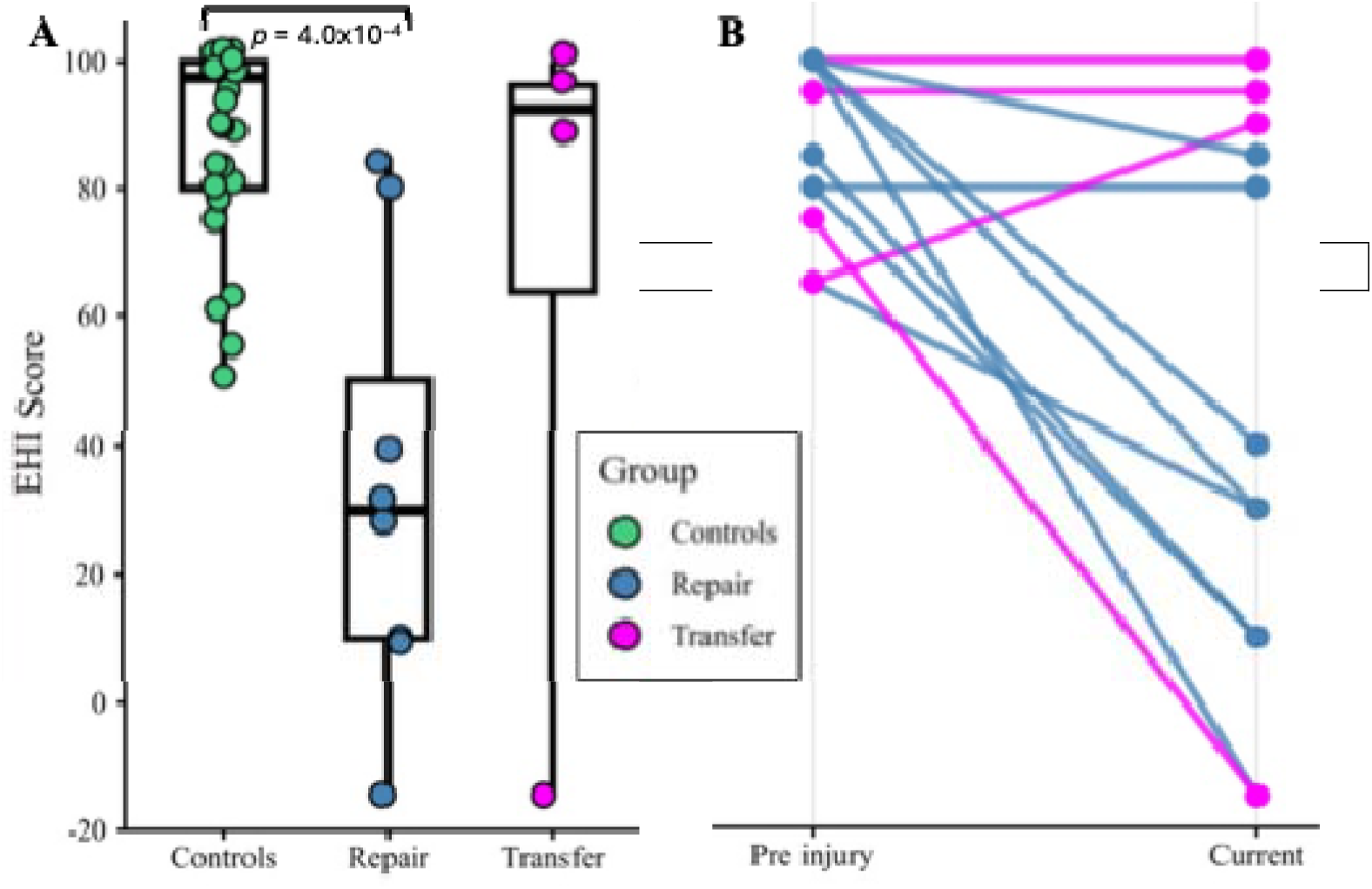
Hand Preference Scores (EHI, Edinburgh Handedness Inventory). Each point represents one participant. **A:** EHI_current_. For patients, “current” = after injury and surgery. **B:** Change between pre-injury and current for individual patients.

EHI_pre-injury_ scores were high and similar in both the Repair group (median 92.5 [65–100]) and the Transfer group (median 85 [65–100]) indicating strong right hand preference.

However, EHI_current_ differed between groups, as shown in **Figure 1A**. Further quantitative analysis (secondary due to small group size) confirmed this finding: EHI_current_ hand preference differed between the three groups (Kruskal-Wallis χ^2^ (2) = 14.62, *p* = 6.7×10^−4^), and post-hoc tests confirmed statistically significant difference between Controls vs. Repair (Z = 3.82, *p-adjusted* = 4.0×10^−4^) but not Controls vs. Transfer (Z = 0.73, *p-adjusted* = 0.4612) or Repair vs. Transfer (Z = −1.82, *p-adjusted* = 0.1351). These results were evident in weaker right hand preference for the Repair group (median 30 [−15–85]) than the Transfer group (median 92.5 [−15–100]), consistent with a pattern in which Transfer participants showed higher right hand preference post-injury and surgery.

Repair patients reported equal or less right hand use after injury and surgery (EHI_current_) compared to before their injury (EHI_pre-injury_), but most transfer patients showed the opposite pattern, as shown in **Figure 1B**. To evaluate within-patient changes in hand preference, hand preference shift scores (ΔEHI = EHI_current_ − EHI_pre-injury_) are reported in **Table 2**. The two groups differed qualitatively, as shown in **Figure 2A**: the Repair group showed reduced right hand preference after injury (ΔEHI median −60 [−115–0]) whereas the Transfer group showed no such reduction, (ΔEHI median 0 [−90 – 25]). In our secondary quantitative analysis, ΔEHI did not differ significantly between groups (W = 8, *p* = 0.1988), but the two groups differed in whether ΔEHI was present: it was present in the Repair group (ΔEHI ≠ 0 via two-tailed signed rank, V = 0, *p* = 0.0222) but not the Transfer group (V = 1, *p* = 1.0), following the same qualitative pattern as above: Transfer but not Repair retained strong right hand preference.

**Figure 2.**
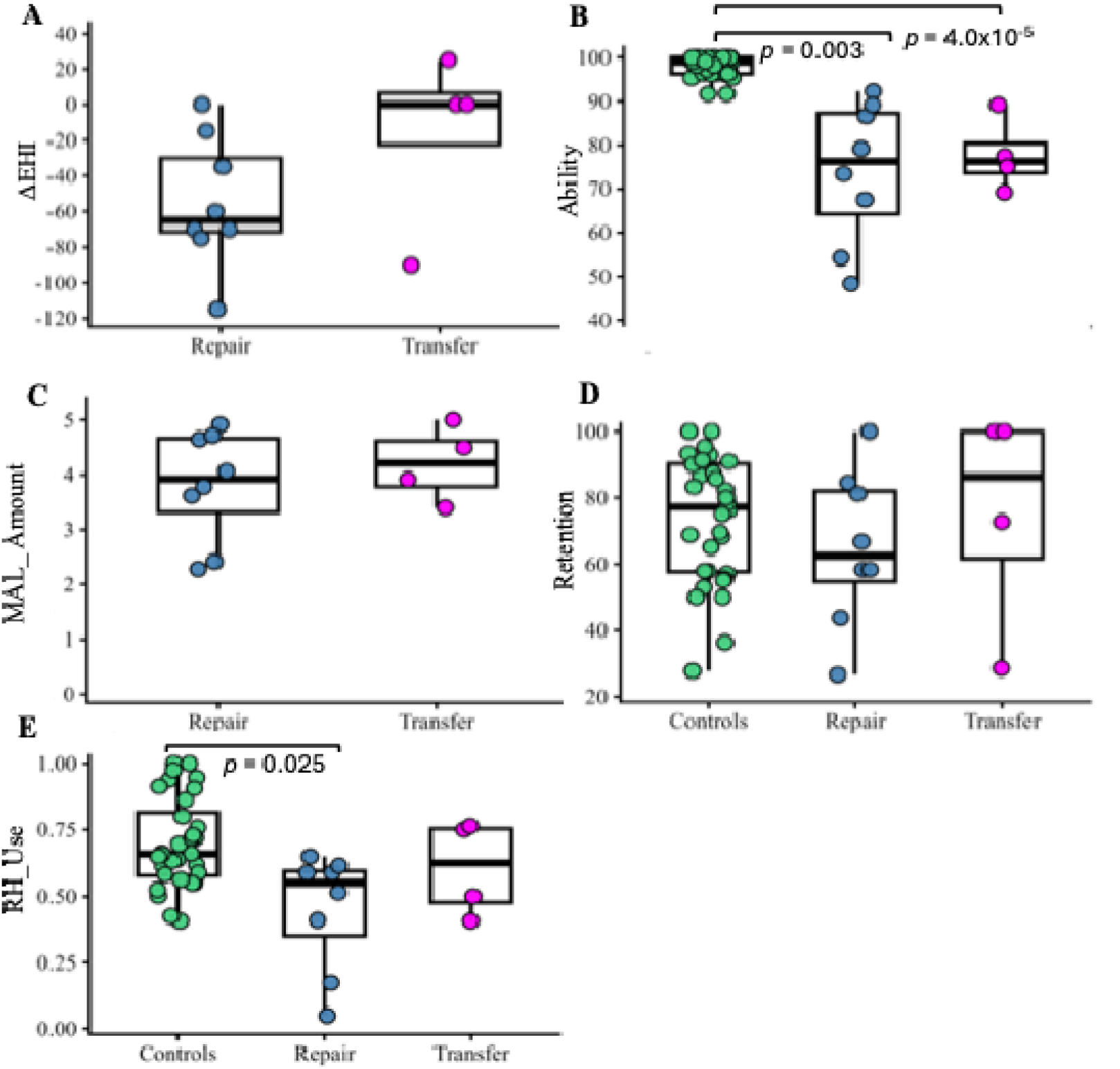
Measures of hand function and use. Boxplots = quartiles, points = individual participants. **A:** Hand preference shift, self-report (ΔEHI). **B:** Upper limb ability, self-report (DASH). **C:** Affected limb use, self-reported (MAL_Amount **D:** Retained participation in precision hand activities, self-report (ACS). **E:** Right hand choices during goal-oriented reach-to-grasp, measured (BBT).

#### Upper-limb ability is significantly lower in patients but did not differ between repair and transfer groups

Upper limb ability (DASH) differed significantly between the three groups (Kruskal-Wallis χ^2^ (2) = 25.64, *p* = 2.71 × 10^−6^), as shown in **Figure 2B**. Post-hoc tests confirmed statistically significant differences between Controls vs. Repair (Z = 4.36, *p-adjusted* = 3.97 × 10^−5^) and Controls vs. Transfer (Z = 3.20, *p-adjusted* = 0.003) but not Repair vs. Transfer (Z = –0.04, *p-adjusted* = 0.968). These differences arose from higher ability for Controls (median 98.9 [91.7–100]) than either Repair (median 76.3 [48.3–92.2]) or Transfer (median 76.3 [69.2–89.2]).

#### Self-reported amount of affected hand use did not differ between repair and transfer groups

Self-reported amount of use of the affected upper limb (MAL_Amount) did not differ between patient groups: the two groups were qualitatively similar, as shown in **Figure 2C**, with overlapping distributions for Repair (median 3.92 [2.28–4.92]) and Transfer (4.21 [3.40–5.00]). Quantitative analysis confirmed this qualitative pattern, with no difference between groups (W= 59, *p* = 0.6711). Of the participants who provided written comments, 6/6 reported ongoing functional limitations including impaired gripping, hesitation, compensation, adaptive equipment, or limited bimanual use; but 4/6 nevertheless reported near-normal to normal use of the affected hand (MAL_Amount 4–5), which points to discordance between MAL_Amount scores and participants’ qualitative descriptions, across participants with varying times since injury and surgery (**Supplementary Table 4)**.

#### Retention of precision-hand activities did not differ between groups

Retention of precision hand activities did not statistically differ between groups (Kruskal-Wallis χ^2^ (2) = 2.41, *p* = 0.3099). Qualitatively, Retention was similar between Controls (77.2% [42.3–100]) and Repair (75.0% [35.6–97.0]), with a trend toward higher Retention in Transfer (83.9% [53.2–98.2]). The broad ranges across groups suggest considerable inter-individual variability in retention as shown in **Figure 2D**.

#### Right hand choices are reduced in repair but relatively preserved in transfer group

Goal-directed right-left hand choices (Block Building Task) differed significantly between the three groups (Kruskal-Wallis χ^2^ (2) = 7.35, *p* = 0.0252), as shown in **Figure 2E**. Post-hoc tests confirmed statistically significant difference between Controls vs. Repair (Z = 2.67, *p-adjusted* = 0.0226) but not Controls vs. Transfer (Z = 0.86, *p-adjusted* = 0.3883) or Repair vs. Transfer (Z = –0.977, *p-adjusted*= 0.6565). This effect was driven by lower RH_Use in the Repair (median 0.55 [0.05–0.65]) compared to the Controls (median 0.66 [0.41–1]) with the Transfer group showing widely distributed values (median 0.63 [0.40–0.76]).

#### Measures of hand function and use were largely independent of each other

In patients, the 5 measures of hand function and use (and time since surgery) were largely uncorrelated, as shown in **Table 3**. The one exception was a significant correlation between ΔEHI and EHI_current_ (ρ = 0.929, FDR-corrected *p* = 1.9×10^−4^) due to the linked nature of two EHI variables.

**Table 3:** Correlations between measures of hand function and use, in patients.

| | Mo. since surgery | RH Use | Retention | MAL Amount | $\Delta\text{EHI}$ | $\text{EHI}_{\text{current}}$ |
| --- | --- | --- | --- | --- | --- | --- |
| Mo. since surgery | - | -0.267 | -0.267 | -0.470 | 0.014 | 0.053 |
| RH Use |  | - | -0.369 | 0.051 | 0.666<sup>†</sup> | 0.539 |
| Retention |  |  | - | 0.106 | -0.167 | -0.076 |
| MAL Amount |  |  |  | - | 0.441 | 0.425 |
| $\Delta\text{EHI}$ | | | | | - | 0.929* |

Note. Values are Spearman's $\rho$ . FDR-corrected $p$ -values: \* = $1.9 \times 10^{-4}$ , <sup>†</sup> = 0.13, others > 0.35.

**Table 4:** Primary hypotheses: clusters with significant connectivity with left M1.

| Cluster name | All cluster regions | Peak region | Peak MNI coordinates | Voxels | Cluster $t$ | Cluster $p$ |
| --- | --- | --- | --- | --- | --- | --- |
| **4.1. Patients > Controls: Figure 3A** |  |  |  |  |  |  |
| Bilateral visual | V1, V2 bilateral; LO left | V1 right | +16, -76, +06 | 2312 | 5.56 | $3.0 \times 10^{-6}$ |
| Bilateral premotor | SFG bilateral | SFG right | +16, +32, +50 | 457 | 4.45 | $6.3 \times 10^{-5}$ |
| **4.2. Transfer > Repair: Figure 3B** |  |  |  |  |  |  |
| Right inferior parietal | IPL, aIPS right | IPL right | +50, -56, +34 | 256 | 7.25 | 0.0246 |
| **4.3. Time since surgery, log-months (voxel threshold $p < 0.05$): Figure 3C** |  |  |  |  |  |  |
| Midline inferior parietal | IPL, precuneus right | IPL right | +44, -68, +30 | 886 | -7.24 | 0.0313 |

*Notes.* Voxel thresholds $p < 0.01$ unless otherwise specified. Clusters 4.2 and 4.3 did not overlap (0%). See text for abbreviations.

### Functional Connectivity with left M1 seed during right hand visuomotor precision drawing

We assessed seed-based functional connectivity (SBC) between left M1 and the rest of the brain during right hand drawing and compared patients with healthy controls, as detailed in the following sections.

#### Patient-specific connectivity between left M1 and visual, premotor, and inferior parietal areas

To test whether patients (Repair, Transfer) would exhibit different task-based FC patterns compared to healthy controls (hypothesis 1), we evaluated between-group differences in SBC with left M1. In the contrast Patients > Controls, patients showed greater SBC with left M1 than controls in two bilateral but asymmetric clusters (**Figure 3A**). The first cluster (“bilateral visual”) had its peak in right primary visual cortex (V1) and extended into bilateral V1, V2, and left lateral occipital cortex (LO). The second cluster (“bilateral premotor”) had its peak in right superior frontal gyrus (SFG) and extended into bilateral SFG, as detailed in **Table 4.1**. The Controls > Patients contrast revealed no significant clusters.

**Figure 3.**
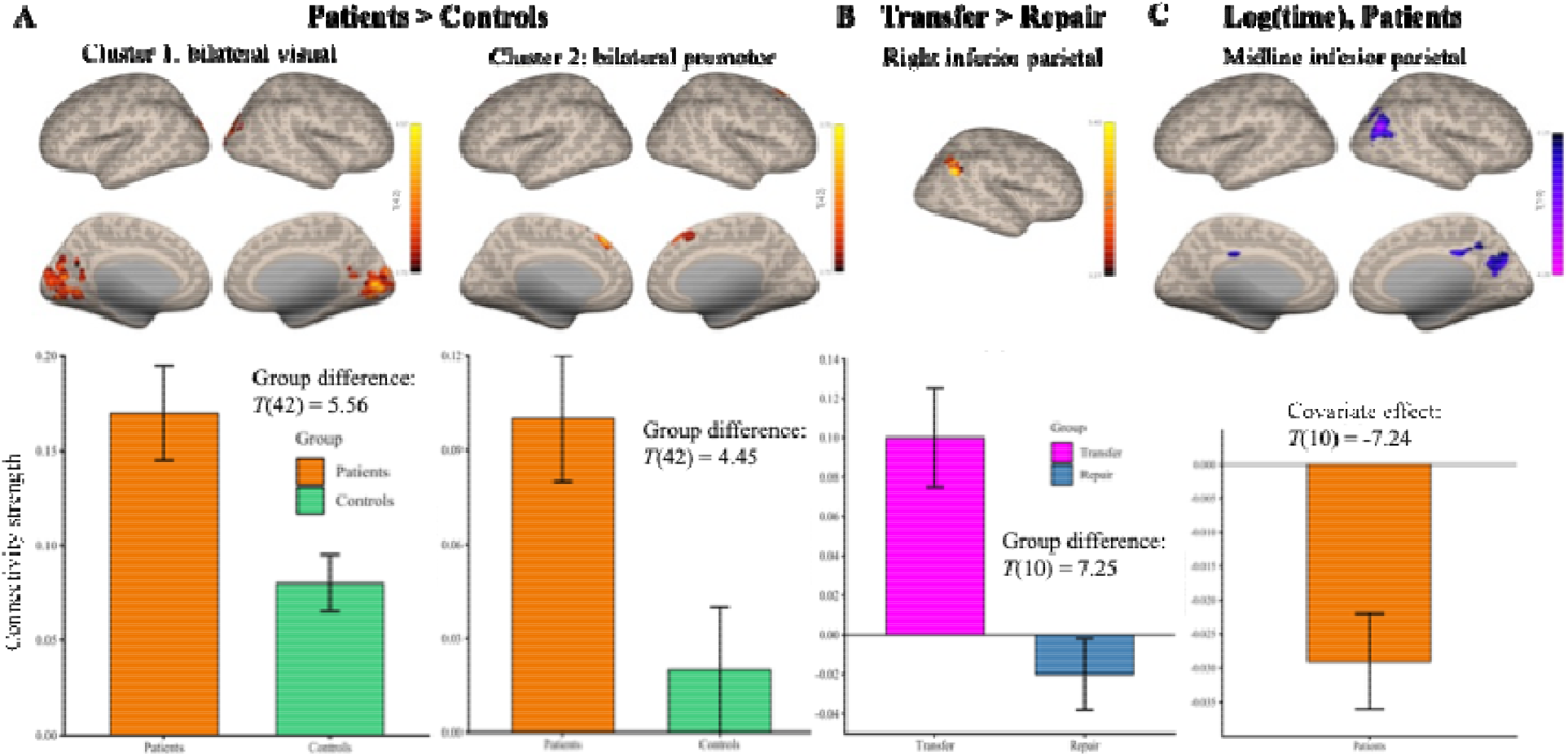
Seed-based functional connectivity with left Ml during right hand drawing *Top row*: Cortical surface maps; voxelwise statistical threshold p < 0.01 unless otherwise specified. *Bottom row*; Cluster connectivity strength (Fisher’s z). mean ± SEM. **A:** Patients > Controls (n = 44). **B:** Transfer > Repair (n = 12). **C:** Log(months) since surgery, patients only in (n = 12). voxelwise threshold *p* < 0.05.

To test whether the Transfer group would exhibit task-based SBC patterns different from the Repair group (hypothesis 2), we evaluated between-group differences in SBC with left M1. In the contrast Transfer > Repair, Transfer patients showed greater SBC with left M1 in a “right inferior parietal” cluster with a peak in right inferior parietal lobule (IPL), extending into right anterior intraparietal sulcus (aIPS) (**Figure 3B** and **Table 4.2**). The contrast Repair > Transfer identified no significant clusters.

In summary, SBC with left M1 during right hand drawing showed distinct clusters that were associated with injury status (Patients vs Controls) and patient groups (Transfer vs Repair), consistent with our first two hypotheses.

#### No linear effect of time since surgery on M1 connectivity, with exploratory evidence of a nonlinear association

To examine whether SBC with left M1 during right hand drawing varied as a function of time, time since surgery was evaluated as a covariate in a group-level SBC analysis of the Patient group (Transfer and Repair, n=12). Time since surgery in months (**Table 2**) was included as a continuous covariate and varied across patients: Repair (median 9 [1-418]) and Transfer (median 16.5 [2-158]). We found no significant clusters, suggesting that FC was not associated with time since surgery.

To explore non-linear relationships between time since surgery and SBC, we performed a post-hoc exploratory analyses of log time and time^2^. No significant clusters were found at *p* < 0.01, but at *p* < 0.05 (uncorrected) revealed one cluster where left M1 FC was negatively associated with log-transformed time since surgery: this “midline inferior parietal” cluster had a peak in the right IPL, extending into right aIPS (**Table 4.3, Figure 3C**). These were the same anatomical regions which were identified in the Transfer > Repair “right inferior parietal” cluster, but the clusters did not overlap (0%). No significant clusters were observed for time^2^ at any threshold.

#### Greater visual and parietal functional connectivity with left M1 in transfer than repair groups relative to healthy controls

To determine whether the Patient > Control differences in left M1 FC were driven by one patient subgroup (i.e. Repair or Transfer), we compared them against the contrasts Repair > Controls and Transfer > Controls. For Repair > Controls we found no significant clusters at our chosen voxel-level threshold (p < 0.01); however, at p < 0.05 (uncorrected), we found two Repair > Control clusters, both of which overlapped heavily with Patient > Control clusters (**Table 5.1, Figure 4A**). For the contrast Transfer > Controls, we found bilateral visual, right inferior parietal, and left visual clusters at our chosen voxel-level threshold (p < 0.01); of these, the visual and parietal clusters overlapped substantially with Patient > Control clusters (**Table 5.2, Figure 4B**). Overall, these results indicate that the FC differences observed in Patients > Controls were not attributable to a single patient group: despite the Transfer group’s stronger and more widespread effects than Repair, the common Patient effects were never group-specific, with substantial overlap between the Patient > Control clusters and subgroup-specific clusters.

**Figure 4.**
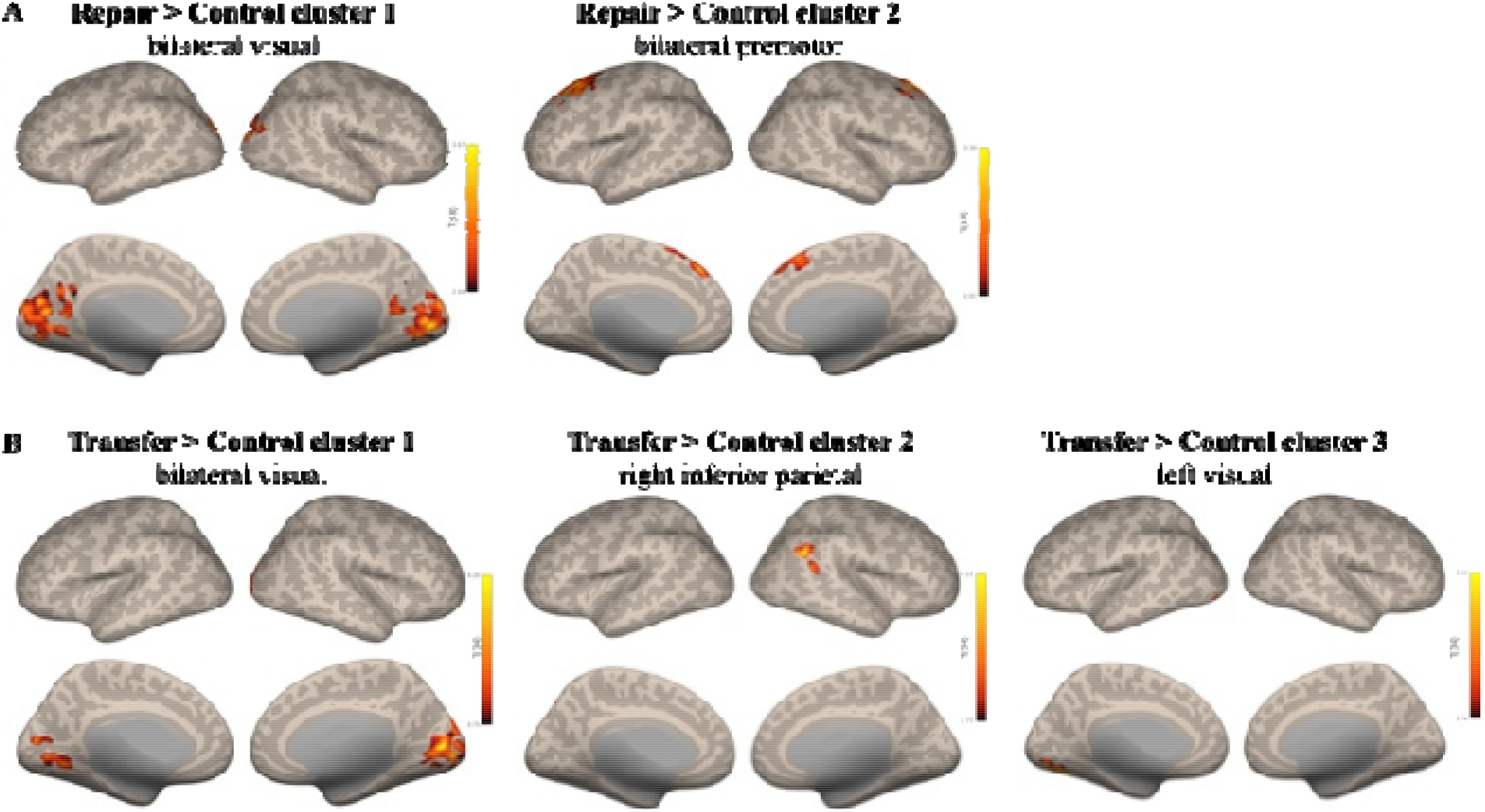
Patient subgroups: cortical surface maps of seed-bused functional connectivity with Ieft MI during right hand drawing. **A:** Repair > Controls, liberal voxelwise statistical threshold (*p* < 0.05). **B:** Transfer > Controls, standard voxelwise statistical threshold (*p* < 0.01).

**Table 5:** Clusters with significant Patient-Subgroup > Control connectivity with left M1.

|  |  |  |  |  |  |  |  |
| --- | --- | --- | --- | --- | --- | --- | --- |
| Cluster name | All cluster regions | Peak region | Peak MNI coordinates | Voxels | Cluster $t$ | Cluster $p$ | Overlap with Pat > Ctl |
| **5.1. Repair > Controls (voxel threshold $p < 0.05$): Figure 4A** |  |  |  |  |  |  |  |
| Midline visual | V1, V2 bilateral | V1 right | +08, -82, +00 | 2375 | 4.66 | $2.0 \times 10^{-5}$ | 79% |
| Bilateral premotor | SFG bilateral | SFG right | +16, +32, +48 | 1700 | 4.15 | 0.0003 | 93% |
| **5.2. Transfer > Controls: Figure 4B** |  |  |  |  |  |  |  |
| Bilateral visual | V1, V2 bilateral | V1 right | +14, -78, +10 | 1437 | 5.51 | $< 1 \times 10^{-6}$ | 66% |
| Right inferior parietal | IPL | IPL right | +52, -52, +36 | 268 | 4.85 | 0.0453 | 43% |
| Left visual | V1, V2, V3V, V4 left | V4 left | -24, -80, -14 | 261 | 4.59 | 0.0453 | 1% |
| *Note.* Voxel thresholds $p < 0.01$ unless otherwise specified. Pat = patient, Ctl = control, Crus = cerebellar crus; see text for other abbreviations. | | | | | | | |

#### Retention and right–left hand choices showed limited association with left M1 FC

We explored whether task-based SBC with left M1 during right-hand drawing was associated with Retention by entering it as a covariate in group-level SBC analysis. No clusters were identified at the predefined threshold (p < 0.01). However, at p < 0.05 (uncorrected), Retention was negatively associated with SBC between left M1 and occipito-cerebellar and temporo-cerebellar areas, and positively associated with SBC between left M1 and midline cingulate areas (**Table 6.1, Figure 5A**, for cerebellar views see **Supplementary Figure 2**). The midline visual cluster overlapped minimally with the bilateral visual cluster for Patients > Controls (14% overlap).

**Figure 5.**
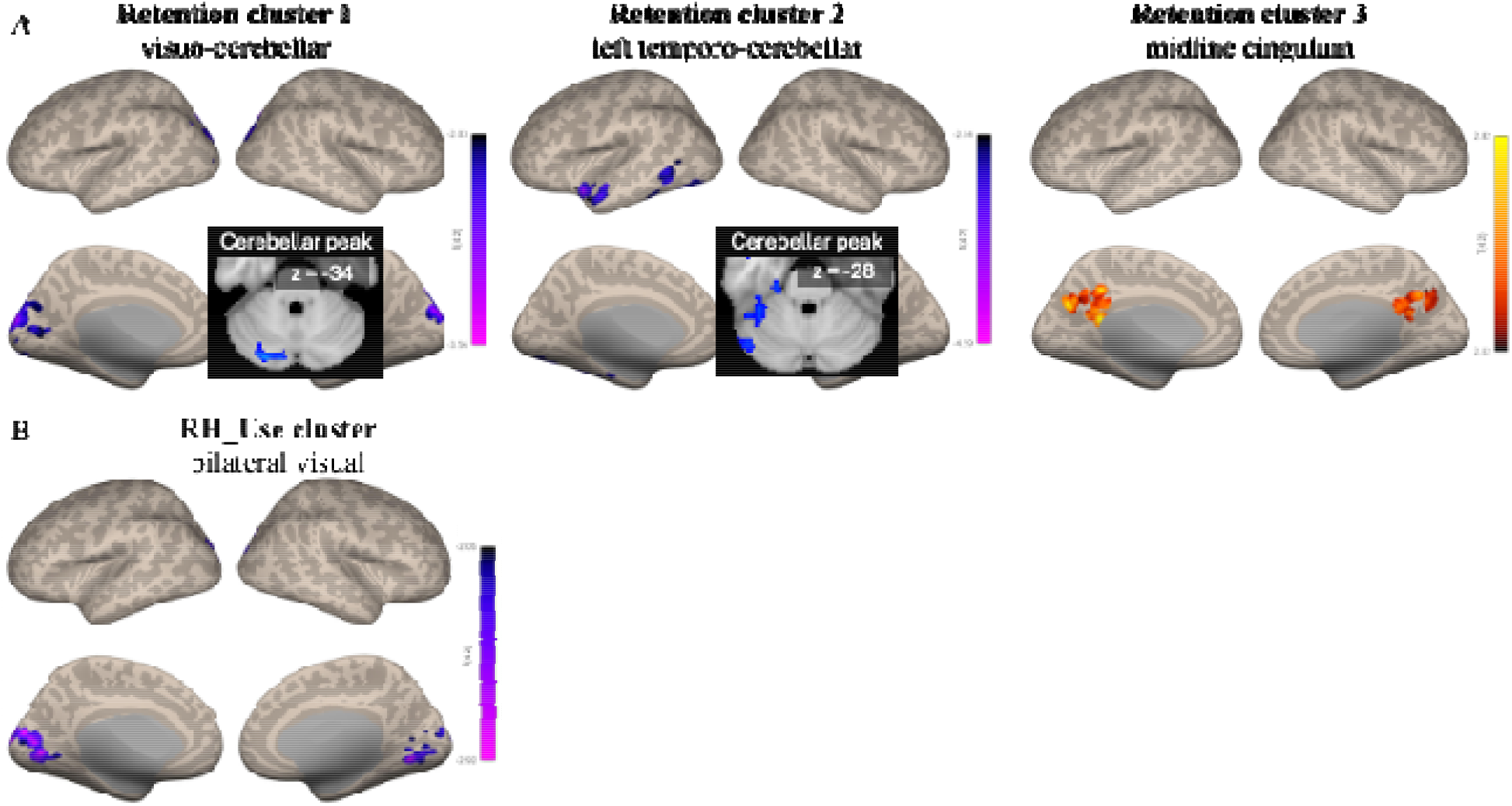
Exploratory analyses: cortical surface maps of seed-based functional connectivity with left Ml during right hand drawing. Liberal voxelwise statistical threshold (*p* < 0 05). Insets = slice view of peak cerebellar voxels, see **Supplementary Figure 2** for cerebellum details. **A:** Retention (Activity Card Sort hand activities). **B:** Right hand use (Block Building Task goal-directed reaches).

Separately, we explored whether task-based SBC with left M1 was associated with the frequency of right-hand choices during goal-directed movement (RH_Use). We again found no clusters at the predefined threshold (p < 0.01). At p < 0.05 (uncorrected), RH_Use was negatively associated with SBC between left M1 and a “bilateral visual” cluster (**Table 6.2, Figure 5B**), which overlapped substantially with the bilateral visual cluster for Patients > Controls (59% overlap).

These analyses were conducted across all participants; no effects were observed in the patient group alone at any threshold. Overall, these exploratory findings suggest a potential trend in which Retention is associated with distributed changes in FC between left M1 and visual-cerebellar and medial cortical regions, while low use of the dominant right hand (across all participants) may involve the same M1-V1 connectivity pattern that distinguished Patients from Controls.

**Table 6:** Exploratory analyses: clusters with significant Retention / Use connectivity with left M1.

| Cluster name | All cluster regions | Peak region | Peak MNI coordinates | Voxels | Cluster $t$ | Cluster $p$ | Overlap with Pat > Ctl |
| --- | --- | --- | --- | --- | --- | --- | --- |
| **6.1. Retention, all participants (voxel threshold $p < 0.05$): Figure 5A** |  |  |  |  |  |  |  |
| Occipito-cerebellar | V1, V2, Crus II left; LO right | LO right | +26, -84, +42 | 2416 | -4.29 | $2.7 \times 10^{-4}$ | 14% |
| Left temporo-cerebellar | Crus I, V4, amygdala, PP, STG left | Crus I left | -38, -04, -18 | 2122 | -6.09 | $5.2 \times 10^{-4}$ | 0% |
| Midline cingulum | Cingulate gyrus left; precuneus midline | White matter (cingulum left) | -10, -48, +20 | 779 | 3.66 | 0.0483 | 0% |
| **6.2. RH_use, all participants (voxel threshold $p < 0.05$): Figure 5B** |  |  |  |  |  |  |  |
| Bilateral visual | V1, V2 bilateral; V3V right | V1 left | -12, -82, +14 | 1076 | -3.53 | 0.0262 | 59% |
| *Note.* Pat = patient, Ctl = control, Crus = cerebellar crus, PP = planum polare; see text for other abbreviations. |  |  |  |  |  |  |  |

## Discussion

This study used a naturalistic precision drawing task to examine task-based functional connectivity (FC) with left M1 during right hand drawing in healthy adults and patients with nerve repair or transfer surgery to the right hand. We used seed-based FC analysis and identified group-related differences in connectivity: patients (vs. healthy controls) showed higher FC with bilateral primary visual and premotor regions, while the Transfer group (vs. Repair group) showed higher FC with right inferior parietal cortex. Exploratory analyses identified a potential negative association between early months post-surgery (i.e. log-transformed time) and FC with a different cluster in right inferior parietal cortex, and that use of the non-dominant left hand in goal-directed reach to grasp may be linked to the same M1-visual FC as found in Patients (vs. Controls).

The effects of surgery and surgical type had inconsistent effects across 5 measures of upper limb function. Self-reported hand preferences and goal-directed right-hand choice showed a similar pattern: both measures differed between the three groups, driven by atypical function in the Repair group, while the Transfer group did not differ from either group. Upper-limb ability was reduced in both patient groups relative to controls. In contrast, self-reported amount of hand use did not differ between patient groups, despite qualitative reports that suggested individualized patterns of adaptation. Finally, retention of precision-hand activities did not differ between the three groups. Overall, these findings indicate that hand function after PNI surgery is not captured by a single dimension.

The patterns observed across the 5 measures of upper limb function are discussed in the next two sections.

### Self report and movement-based measures reveal different group level patterns

Among the four self-report measures of upper limb function, only the DASH_Ability distinguished patients from healthy controls. Group-level differences (22 points) exceeded the DASH minimal clinically important difference (~10 points) (Gummesson et al., 2003). In contrast, retention of precision-hand activities (i.e., continued participation in such activities outside the clinic) did not differ between patients and controls or between patient groups. However, retention values were widely distributed across all groups, with the greatest variability observed in controls. This substantial within-group variability may have obscured group-level differences and suggests that participation in precision activities is not well-determined by motor impairment. Consistent with this interpretation, prior work suggested that motor activities retention following PNI is influenced by compensatory strategies and psychosocial factors (Gassass, Zhou, et al., 2025).

Goal-directed right–left hand choices revealed a different group pattern. Unlike self-report measures, the BBT assesses (directly, without self-report) a quantitative measure of how often individuals choose to use each hand for goal-directed reach-to-grasp in a context where they can accomplish the reaches with either limb (Kim et al., 2025). At the group level, right (affected) hand choices were less frequent in patients than healthy controls, with this difference driven primarily by low right (affected) hand use in the Repair group. This pattern suggests patients’ behavioral tendency to avoid use of the affected limb despite their continued ability to perform the task with either hand: only 1/12 patients used their unaffected hand on 100% of reaches. Notably, this objective pattern mirrored the Repair group’s shift in self-reported hand preference, indicating that the shift reflects a true group level change in limb use tendency rather than a measurement artifact. Similarly, previous studies have suggested that functional hand use after PNI reflects compensatory strategies rather than motor recovery alone (Chemnitz et al., 2013). Furthermore, right hand choices were not associated with time since surgery. These findings suggest that reduced hand use does not simply reflect recovery stage but may instead represent persistent behavioral adaptation.

### Repair and transfer groups demonstrated individualized patterns of adaptation

Repair and Transfer patients responded differently to self-reported assessments of hand use. This likely reflects measurement and sampling factors, as self-report captures perceived ability rather than motor strategy (Novak et al., 2009) and group comparisons are complicated by individual variability in daily function and time since surgery (**Table 2**). In this context, group-level comparisons may not capture meaningful individual differences.

Consistent with the above interpretation, inter-individual variability was high for all measures (**Figure 2**; **Table 2**). Several individuals in the Repair group exhibited decreased preference for the right hand (EHI) after injury and surgery (**Figure 1**). However, ΔEHI scores varied independently of DASH_Ability, MAL_Amount and time since surgery (**Table 3**).

Despite the measures of function’s high inter-individual variability and lack of consistent group-related effects, the FC results showed distinct group-related effects, as described in the next section.

## Functional Connectivity

### Seed-based connectivity with left M1 identified spatially distinct clusters in patients during right-hand drawing

Our first hypothesis was supported: patients showed distinct FC pattern compared to Controls. Specifically, patients (Repair, Transfer) showed greater task-based connectivity between left M1 and two clusters (**Table 4.1**).

The first patient-specific cluster was centered in primary visual cortex (V1), which encodes retinotopic visual information and edge features with high spatial precision (Wandell et al., 2007; Wu et al., 2012). This level of visual processing is essential for the drawing task which requires continuous monitoring of pen position. Also, accurate task performance depends on the integration of visual and proprioceptive input (Ernst & Banks, 2002; Proske & Gandevia, 2012). However, the likely altered proprioceptive input in PNI patients (Ernst, 2022; Navarro et al., 2007) may increase reliance on visual feedback. Even over months to years, some proprioceptive input returns via regenerated afferents, but because central synaptic connections are reduced, patients often have persistent abnormalities in fine motor coordination (Duff, 2005). Therefore, the increased V1-M1 FC in patients could represent adaptive neural patterns to support increased visual (as opposed to proprioceptive) guidance over motor control.

The second patient-specific cluster was centered in bilateral premotor area, predominantly ipsilateral to the drawing hand. The premotor cortex plays a critical role in higher order motor planning particularly under cognitive demand or reduced movement automaticity (Hoshi & Tanji, 2007; Picard & Strick, 2001). This is especially relevant following PNI surgery, where motor output must be relearned under decreased and altered afferent input (Lundborg, 2000; Taylor et al., 2009) which may also contribute to the ipsilateral dominance connectivity pattern observed here. Our results are consistent with previous studies that demonstrated recruitment of ipsilateral motor and frontal regions during thumb–index tapping of the affected limb after brachial plexus nerve transfer (Dimou et al., 2013).

Our second hypothesis was also supported: FC differed between Transfer and Repair groups. Specifically, Transfer group showed increased FC between left M1 and two clusters: in the right inferior parietal lobule (IPL) and right anterior intraparietal sulcus (aIPS). These parietal regions are central to visuomotor transformation and updating of sensorimotor mappings (Andersen & Cui, 2009; Culham & Valyear, 2006; Rolls et al., 2023). In nerve transfer surgery, a donor nerve is rerouted to reinnervate a new motor target (Moore & Novak, 2014), which demands cortical adaptation between the original donor nerve and the recipient muscle representation (Anastakis et al., 2008). Increased M1–parietal FC in Transfer group may therefore reflect enhanced reliance on parietal mechanisms to stabilize motor output. Interpretations remain cautious given the small Transfer group sample size (n = 4).

### No linear effect of time since surgery on M1 connectivity, but exploratory evidence for a nonlinear effect

The third hypothesis that FC with left M1would covary with time since surgery across all patients was not supported. However, our exploratory analysis showed a negative association between log-transformed time and left M1 FC in the right/medial inferior parietal areas (distinct from the the right IPL cluster in the Patient > Control analysis). A log-time relationship suggests diminishing time effects: larger changes occurring early after surgery and smaller incremental changes later. However, this logarithmic pattern does not indicate complete recovery in this sample but may reflect gradual adaptation and reduced reliance on parietal visuomotor integration as time progressed (Dayan & Cohen, 2011) even as recovery remain incomplete. These exploratory findings should be interpreted cautiously given the small sample size and the use of a more liberal statistical threshold (p < 0.05).

### Transfer vs. Repair: Greater visual and parietal functional connectivity with left M1 in transfer than repair groups relative to healthy controls

To determine whether one patient group drove the Patient > Control differences in SBC with left M1, we conducted SBC analyses to compare Repair and Transfer patients with controls during right-hand drawing. No effects were observed at the predefined threshold (p < 0.01); findings emerged only at p < 0.05 (uncorrected) across all participants and are therefore considered exploratory. Nevertheless, Repair > Control and Transfer > Control effects largely agreed with the Patient > Control effects, especially in V1 and IPL clusters. The Transfer > Control effects were stronger than the Repair > Control effects, which could reflect spurious relationships due to the small Transfer sample size, or may reflect a distinct feature of Transfer patients. If the latter, Transfer patients’ more-atypical patterns may arise because nerve transfer imposes greater demands on cortical adaptation, as rerouted efferent pathways require establishment of novel sensorimotor mappings (Shen, 2022; Yu et al., 2017). In contrast, nerve repair is frequently associated with misdirected axonal reinnervation (Gordon, 2020) which can degrade the quality of sensory feedback that reaches the cortex. Consistent with the reduced affected-hand use observed in the Repair group, their weaker FC effects likely reflect reduced engagement of affected-limb networks rather than complete functional reintegration (Nudo, 2013).

### Activity retention and right-hand choices associated with reduced reliance on vision

We explored the relationship between drawing-related FC with left M1, and life-relevant measures of activity retention and goal-directed hand use. For both, we found significant FC only at a liberal statistical threshold and when all participants were analyzed together. These findings should therefore be interpreted cautiously and viewed as exploratory rather than confirmatory.

Activity retention was negatively associated with FC between left M1 and occipital and cerebellar regions, which may suggest that greater retention of precision hand activities is accompanied by reduced reliance on visually guided control. Activity retention was positively associated with a cluster in the precuneus, cingulate gyrus, adjacent white-matter regions that correspond to the cingulum, and callosal body; which may reflect regionally specific association in which higher activity retention is linked to greater engagement of regions involved in sensorimotor integration, attention, and self-generated action (Cavanna & Trimble, 2006; Lundborg, 2000).

More frequent use of the dominant right hand was negatively associated with FC between left M1 and a cluster in bilateral (right-lateralized) visual cortex, which overlapped heavily with the cluster in bilateral (right-lateralized) visual cortex associated with the Patient > Control group effect. In other words, FC between left M1 and V1 was greater for patients, and also for individuals (patient or otherwise) who used their dominant right hand less often. This pattern may reflect reduced dependence on visual processing with more typical hand use in right-handed adults, though it is difficult to interpret the value of high or low use of the dominant hand: mild-to-moderate PNI can have unpredictable effects on left/right hand choices (Kim et al., 2025; Philip et al., 2022), and in healthy adults, use of the non-dominant hand can reflect adaptive flexibility (Prichard et al., 2013). Regardless, the common pattern of left M1-bilateral V1 FC appeared across different environments (during fMRI vs. outside fMRI), contexts (constrained vs. goal-directed action), and visually guided dexterous tasks (precision drawing vs. reach-to-grasp), which suggests that this pattern represents a consistent, behaviorally relevant mechanism.

### Directions for Future Research

The current study findings point to four important future research directions. *First*, task-based FC should be examined at longitudinal time points in a within-subject design to determine whether the observed visual, premotor, and parietal connectivity effects are consistent across patients or are driven by individual adaptation profiles. *Second*, regions identified in the present study should be used as seed areas to establish whether the observed effects are specific to M1 connectivity or reflect broader network-level adaptations. While the present analyses focused on left M1 as the seed region, future work should capture more distributed changes across the sensorimotor system (Gassass, 2025). *Third*, future studies should assess whether individual differences in connectivity are associated with drawing performance, symptom severity, and neurophysiological measures of peripheral regeneration. This would help determine whether clinical and neurophysiological measures reflect underlying constraints on cortical plasticity. *Fourth*, to clarify how individual connectivity patterns relate to behavior and recovery, fMRI analytical methods should move beyond group-averaged results to methods that capture inter-individual variability and within-individual FC.

### Limitations

The study has several limitations. *First*, the cross-sectional design limits inference regarding the temporal evolution of connectivity changes after PNI surgery. However, this study represents the first investigation of task-based functional connectivity during an ecologically valid, dexterous hand task in this population. *Second*, the sample size was modest and clinically heterogeneous. Although this reflects the rarity of the population, heterogeneity may have reduced sensitivity to detect consistent effects and limits generalizability. *Third*, the inclusion criteria required participants to have sufficient motor function to complete the precision drawing task, which may have biased the sample toward higher-functioning individuals and limits generalizability to patients with more severe impairment. Nevertheless, qualitative measures indicated persistent patterns of adaptation accompanied by substantial functional deficits. *Fourth*, several connectivity effects observed only at p < 0.05 (uncorrected) and should be interpreted as exploratory. However, their spatial overlap with visual and higher-order control networks suggests potential relevance for compensatory motor processes and future hypothesis generation. Because this threshold was applied at the voxel rather than cluster level, it primarily affects cluster extent rather than anatomical localization (Woo et al., 2014).

## Conclusion

We used a dexterous ecologically valid, visuospatial fine motor task during fMRI to capture behaviorally relevant patterns of functional connectivity in patients after peripheral nerve injury (PNI) and surgery to the dominant right upper limb. Task-based functional connectivity revealed a consistent pattern of compensation despite clinical heterogeneity and inconsistent patterns across measures of upper limb function and use. During a visuomotor right hand drawing task, patients with right hand PNI surgery (compared to healthy adults) showed greater functional connectivity between the affected left M1 and bilateral visual and premotor regions. This pattern is consistent with greater reliance on vision and input from higher order regions with increased cognitive effort and when sensorimotor feedback is compromised, a pattern known to result from PNI.

Patients with nerve transfer surgery, compared to repair surgery, exhibited greater connectivity between left M1 and right inferior parietal lobule, possibly due to additional visuomotor demands required to integrate novel efferent sensorimotor pathways. Exploratory results suggests that in early months after surgery, patients initially depend heavily on parietal visuospatial monitoring, and this dependence decreases as motor patterns stabilize (even imperfectly). Exploratory results also suggest that high reliance on vision (as found in PNI patients) is associated with low use of the dominant hand in all individuals.

These findings have important clinical implications. *First*, rehabilitation after PNI surgery should incorporate early, functionally relevant, and appropriately challenging activities to engage the brain regions that support task performance. *Second*, distinct connectivity patterns across patient groups and across time since surgery suggest that task-related connectivity varies with surgical and recovery context which support the need to tailor rehabilitation in terms of task selection and progression across stages of recovery. *Finally*, the lack of consistent patterns across self-report and task-based measures highlights that hand function after PNI surgery is multidimensional. Rehabilitation assessment approaches should aim to capture impairment, real-life hand use, and performance-based outcomes to plan targeted interventions.

## Data availability statement

The data that support the findings of this study will be made openly available upon completion of clinical trial NCT05207878.

## References

Amini, A., Fischmeister, F. P. S., Matt, E., Schmidhammer, R., Rattay, F., & Beisteiner, R. (2018). Peripheral nervous system reconstruction reroutes cortical motor output—Brain reorganization uncovered by effective connectivity. Frontiers in Neurology, 9, 420883.

Amunts, K., Mohlberg, H., Bludau, S., & Zilles, K. (2020). Julich-Brain: A 3D probabilistic atlas of the human brain’s cytoarchitecture. Science, 369(6506), 988–992.

Anastakis, D. J., Malessy, M. J., Chen, R., Davis, K. D., & Mikulis, D. (2008). Cortical plasticity following nerve transfer in the upper extremity. Hand clinics, 24(4), 425–444.

Andersen, R. A., & Cui, H. (2009). Intention, action planning, and decision making in parietal-frontal circuits. Neuron, 63(5), 568–583.

Anthony, M., Hattori, R., Nicholas, M. L., Randolph, S., Lee, Y., Baum, C. M., & Connor, L. T. (2024). Social Support Mediates the Association between Abilities and Participation after Stroke. OTJR: Occupational Therapy Journal of Research, 15394492241249446.

Avants, B. B., Tustison, N. J., Song, G., Cook, P. A., Klein, A., & Gee, J. C. (2011). A reproducible evaluation of ANTs similarity metric performance in brain image registration. NeuroImage, 54(3), 2033–2044. 10.1016/j.neuroimage.2010.09.025

Baum, C. M., & Edwards, D. (2008). ACS: activity card sort:Activity card sort: test manual. (No Title).

Beaton, D. E., Davis, A. M., Hudak, P., & McConnell, S. (2001). The DASH (Disabilities of the Arm, Shoulder and Hand) outcome measure: what do we know about it now? The British Journal of Hand Therapy, 6(4), 109–118.

Beaulieu, J. Y., Blustajn, J., Teboul, F., Baud, P., De Schonen, S., Thiebaud, J. B., & Oberlin, C. (2006). Cerebral plasticity in crossed C7 grafts of the brachial plexus: an fMRI study. Microsurgery: Official Journal of the International Microsurgical Society and the European Federation of Societies for Microsurgery, 26(4), 303–310.

Behzadi, Y., Restom, K., Liau, J., & Liu, T. T. (2007). A component based noise correction method (CompCor) for BOLD and perfusion based fMRI. NeuroImage, 37(1), 90–101.

Beisteiner, R., Höllinger, I., Rath, J., Wurnig, M., Hilbert, M., Klinger, N., Geißler, A., Fischmeister, F., Wöber, C., & Klösch, G. (2011). New type of cortical neuroplasticity after nerve repair in brachial plexus lesions. Archives of neurology, 68(11), 1467–1470.

Bhat, D. I., Indira Devi, B., Bharti, K., & Panda, R. (2017). Cortical plasticity after brachial plexus injury and repair: a resting-state functional MRI study. Neurosurg Focus, 42(3), E14. 10.3171/2016.12.FOCUS16430

Cavanna, A. E., & Trimble, M. R. (2006). The precuneus: a review of its functional anatomy and behavioural correlates. Brain, 129(3), 564–583. 10.1093/brain/awl004

Chai, X. J., Castañón, A. N., Öngür, D., & Whitfield-Gabrieli, S. (2012). Anticorrelations in resting state networks without global signal regression. Neuroimage, 59(2), 1420–1428.

Chemnitz, A., Dahlin, L. B., & Carlsson, I. K. (2013). Consequences and adaptation in daily life– patients’ experiences three decades after a nerve injury sustained in adolescence. BMC musculoskeletal disorders, 14(1), 252.

Chen, A., Yao, J., Kuiken, T., & Dewald, J. P. A. (2013). Cortical motor activity and reorganization following upper-limb amputation and subsequent targeted reinnervation. Neuroimage: clinical, 3, 498–506.

Chumbley, J., Worsley, K., Flandin, G., & Friston, K. (2010). Topological FDR for neuroimaging. NeuroImage, 49(4), 3057–3064. 10.1016/j.neuroimage.2009.10.090

Cramer, S. C., Sur, M., Dobkin, B. H., O’Brien, C., Sanger, T. D., Trojanowski, J. Q., Rumsey, J. M., Hicks, R., Cameron, J., & Chen, D. (2011). Harnessing neuroplasticity for clinical applications. Brain, 134(6), 1591–1609.

Culham, J. C., & Valyear, K. F. (2006). Human parietal cortex in action. Current opinion in neurobiology, 16(2), 205–212.

Cutkosky, M. R. (1989). On grasp choice, grasp models, and the design of hands for manufacturing tasks. IEEE Transactions on robotics and automation, 5(3), 269–279.

Dayan, E., & Cohen, L. G. (2011). Neuroplasticity subserving motor skill learning. Neuron, 72(3), 443–454.

Dimou, S., Biggs, M., Tonkin, M., Hickie, I. B., & Lagopoulos, J. (2013). Motor cortex neuroplasticity following brachial plexus transfer. Front Hum Neurosci, 7, 500. 10.3389/fnhum.2013.00500

Duff, S. V. (2005). Impact of peripheral nerve injury on sensorimotor control. J Hand Ther, 18(2), 277–291. 10.1197/j.jht.2005.02.007

Eickhoff, S. B., Paus, T., Caspers, S., Grosbras, M.-H., Evans, A. C., Zilles, K., & Amunts, K. (2007). Assignment of functional activations to probabilistic cytoarchitectonic areas revisited. Neuroimage, 36(3), 511–521.

Ernst, J. W. T.,; Wanke, N.; Frahm, J.; Felmerer, G.; Farina, D.; Schilling, A. F.; Wilke, M. A. (2022). Case Report: Plasticity in Central Sensory Finger Representation and Touch Perception After Microsurgical Reconstruction of Infraclavicular Brachial Plexus Injury [Article]. Frontiers in Neuroscience, 16. 10.3389/fnins.2022.793036

Ernst, M. O., & Banks, M. S. (2002). Humans integrate visual and haptic information in a statistically optimal fashion. Nature, 415(6870), 429–433.

Fischl, B., Salat, D. H., Busa, E., Albert, M., Dieterich, M., Haselgrove, C., Van Der Kouwe, A., Killiany, R., Kennedy, D., & Klaveness, S. (2002). Whole brain segmentation: automated labeling of neuroanatomical structures in the human brain. Neuron, 33(3), 341–355.

Fischmeister, F. P. S., Amini, A., Matt, E., Reinecke, R., Schmidhammer, R., & Beisteiner, R. (2020). A New Rehabilitative Mechanism in Primary Motor Cortex After Peripheral Trauma. Front Neurol, 11, 125. 10.3389/fneur.2020.00125

Fischmeister, F. P. S., Amini, A., Matt, E., Reinecke, R., Schmidhammer, R., & Beisteiner, R. (2020). A new rehabilitative mechanism in primary motor cortex after peripheral trauma. Frontiers in Neurology, 11, 125.

Fraiman, D., Miranda, M. F., Erthal, F., Buur, P., Elschot, M., Souza, L., Rombouts, S., Schimmelpenninck, C., Norris, D., & Malessy, M. (2016). Reduced functional connectivity within the primary motor cortex of patients with brachial plexus injury. Neuroimage: clinical, 12, 277–284.

Friston, K. J., Williams, S., Howard, R., Frackowiak, R. S., & Turner, R. (1996). MovementLrelated effects in fMRI timeLseries. Magnetic Resonance in Medicine, 35(3), 346–355.

Gassass, S., Lipsey, K., Ratner, A. M., & Philip, B. A. (2025). Cortical Plasticity Following Nerve Transfers in the Upper Extremity: A Systematic Review. Washington University School of Medicine in St. Louis.

Gassass, S., Zhou, R., Hattori, R., Liu, L., Connor, L. T., & Philip, B. A. (2025). Activity Loss and Retention Patterns Following Unilateral Peripheral Nerve Injury: Implications for Rehabilitation. The American Journal of Occupational Therapy, 79(6), 7906205100.

Gassass, S. L., Kim; Ratner, Allison M.; Philip Benjamin A. (2025). Cortical Plasticity following Nerve Transfers in the Upper Extremity: A Systematic Review. Program in Occupational Therapy, Washington University School of Medicine in St. Louis, St. Louis, MO, USA Bernard Becker Medical Library, Washington University School of Medicine in St. Louis, St. Louis, MO, USA.

Gorgolewski, K., Burns, C. D., Madison, C., Clark, D., Halchenko, Y. O., Waskom, M. L., & Ghosh, S. S. (2011). Nipype: a flexible, lightweight and extensible neuroimaging data processing framework in python. Frontiers in neuroinformatics, 5, 13. 10.3389/fninf.2011.00013

Groppe, D. (2015). fdr_bh. In (Version 2.3.0.0) MATLAB Central File Exchange.

David Groppe (2026). fdr_bh (https://www.mathworks.com/matlabcentral/fileexchange/27418-fdr_bh), MATLAB Central File Exchange. Retrieved April 15, 2026.

Gummesson, C., Atroshi, I., & Ekdahl, C. (2003). The disabilities of the arm, shoulder and hand (DASH) outcome questionnaire: longitudinal construct validity and measuring self-rated health change after surgery. BMC musculoskeletal disorders, 4(1), 11.

Hallquist, M. N., Hwang, K., & Luna, B. (2013). The nuisance of nuisance regression: spectral misspecification in a common approach to resting-state fMRI preprocessing reintroduces noise and obscures functional connectivity. Neuroimage, 82, 208–225.

Hoshi, E., & Tanji, J. (2007). Distinctions between dorsal and ventral premotor areas: anatomical connectivity and functional properties. Current opinion in neurobiology, 17(2), 234–242.

Hudak, P. L., Amadio, P. C., Bombardier, C., Beaton, D., Cole, D., Davis, A., Hawker, G., Katz, J. N., Makela, M., & Marx, R. G. (1996). Development of an upper extremity outcome measure: the DASH (disabilities of the arm, shoulder, and head). American journal of industrial medicine, 29(6), 602–608.

Isaacs, J. (2013). Major peripheral nerve injuries. Hand clinics, 29(3), 371–382.

Jenkinson, M., Bannister, P., Brady, M., & Smith, S. (2002). Improved optimization for the robust and accurate linear registration and motion correction of brain images. NeuroImage, 17(2), 825–841.

Kaas, J. H. (2000). The reorganization of somatosensory and motor cortex after peripheral nerve or spinal cord injury in primates. Progress in brain research, 128, 173–179.

Kapil, N., Kim, T., Gassass, S., Zhou, R., Carter, A., Dobbins, I., Liu, L., McAvoy, M., Wheelock, M., Wang, Y., Brogan, D., Dy, C., Mackinnon, S., & Philip, B. (2025). Neural mechanisms of handedness for precision drawing: hand-dependent engagement of cortical networks for bimanual control and tool use. bioRxiv, 2025.2011.2018.689091. 10.1101/2025.11.18.689091

Katz, N., Karpin, H., Lak, A., Furman, T., & Hartman-Maeir, A. (2003). Participation in occupational performance: Reliability and validity of the Activity Card Sort. OTJR: Occupation, Participation and Health, 23(1), 10–17.

Khan, H., & Perera, N. (2020). Peripheral nerve injury: an update. Orthopaedics and Trauma, 34(3), 168–173.

Kim, T., Fletcher, S., Gonzalez, C. L. R., & Philip, B. A. (2025). Block Building Task Identifies Distinct Groups of Left/Right-hand Choice Patterns After Unilateral Peripheral Nerve Injury. J Vis Exp(217). 10.3791/66919

Kim, T., Zhou, R., Gassass, S., Soberano, T., Liu, L., & Philip, B. A. (2024). Healthy adults favor stable left/right hand choices over performance at an unconstrained reach-to-grasp task. Experimental brain research. 10.1007/s00221-024-06828-5

Kline, D. G. (1990). Surgical repair of peripheral nerve injury. Muscle & Nerve: Official Journal of the American Association of Electrodiagnostic Medicine, 13(9), 843–852.

Lee, S. K., & Wolfe, S. W. (2000). Peripheral nerve injury and repair. JAAOS-Journal of the American Academy of Orthopaedic Surgeons, 8(4), 243–252.

Lemon, R. N. (2008). Descending pathways in motor control. Annu. Rev. Neurosci., 31(1), 195–218.

Li, T., Hua, X. Y., Zheng, M. X., Wang, W. W., Xu, J. G., Gu, Y. D., & Xu, W. D. (2015). Different cerebral plasticity of intrinsic and extrinsic hand muscles after peripheral neurotization in a patient with brachial plexus injury: A TMS and fMRI study. Neurosci Lett, 604, 140–144. 10.1016/j.neulet.2015.07.015

Lundborg, G. (2000). Brain plasticity and hand surgery: an overview. The Journal of Hand Surgery: British & European Volume, 25(3), 242–252.

Machado, S., Cunha, M., Velasques, B., Minc, D., Teixeira, S., Domingues, C. A., Silva, J. G., Bastos, V. H., Budde, H., & Cagy, M. (2010). Sensorimotor integration: basic concepts, abnormalities related to movement disorders and sensorimotor training-induced cortical reorganization. Rev Neurol, 51(7), 427–436.

Mathiowetz, V., Volland, G., Kashman, N., & Weber, K. (1985). Adult norms for the Box and Block Test of manual dexterity. The American Journal of Occupational Therapy, 39(6), 386–391.

Meyers, E. C., Kasliwal, N., Solorzano, B. R., Lai, E., Bendale, G., Berry, A., Ganzer, P. D., Romero-Ortega, M., Rennaker, R. L., & Kilgard, M. P. (2019). Enhancing plasticity in central networks improves motor and sensory recovery after nerve damage. Nature communications, 10(1), 5782.

Moore, A. M. (2014). Nerve transfers to restore upper extremity function: a paradigm shift. Frontiers in Neurology, 5, 40.

Moore, A. M., & Novak, C. B. (2014). Advances in nerve transfer surgery. J Hand Ther, 27(2), 96–104; quiz 105. 10.1016/j.jht.2013.12.007

Napier, J. R. (1956). The prehensile movements of the human hand. The Journal of Bone & Joint Surgery British Volume, 38(4), 902–913.

Navarro, X. (2009). Neural plasticity after nerve injury and regeneration. International review of neurobiology, 87, 483–505.

Navarro, X., Vivó, M., & Valero-Cabré, A. (2007). Neural plasticity after peripheral nerve injury and regeneration. Progress in Neurobiology, 82(4), 163–201.

Nieto-Castanon, A. (2020). Handbook of functional connectivity Magnetic Resonance Imaging methods in CONN. 10.56441/hilbertpress.2207.6598

Nieto-Castanon, A. (2025). Preparing fMRI data for statistical analysis. In fMRI techniques and protocols (pp. 163–191). Springer.

Nieto-Castanon, A., & Whitfield-Gabrieli, S. (2021). CONN functional connectivity toolbox (RRID: SCR_009550), Version 21. Series CONN functional connectivity toolbox (RRID: SCR_009550), Version, 21.

Nordmark, P. F., Ljungberg, C., & Johansson, R. S. (2018). Structural changes in hand related cortical areas after median nerve injury and repair. Scientific reports, 8(1), 4485.

Novak, C. B., Anastakis, D. J., Beaton, D. E., & Katz, J. (2009). Patient-reported outcome after peripheral nerve injury. The Journal of hand surgery, 34(2), 281–287.

Nudo, R. J. (2013). Recovery after brain injury: mechanisms and principles. Frontiers in Human Neuroscience, 7, 887.

Oldfield, R. C. (1971). The assessment and analysis of handedness: the Edinburgh inventory. Neuropsychologia, 9(1), 97–113.

Patriat, R., Reynolds, R. C., & Birn, R. M. (2017). An improved model of motion-related signal changes in fMRI. NeuroImage, 144, 74–82.

Pereira, C. T., Hill, E. E., Stasyuk, A., Parikh, N., Dhillon, J., Wang, A., & Li, A. (2023). Molecular basis of surgical coaptation techniques in peripheral nerve injuries. Journal of Clinical Medicine, 12(4), 1555.

Philip, B. A., & Frey, S. H. (2014). Compensatory changes accompanying chronic forced use of the nondominant hand by unilateral amputees. Journal of Neuroscience, 34(10), 3622–3631.

Philip, B. A., & Frey, S. H. (2016). Increased functional connectivity between cortical hand areas and praxis network associated with training-related improvements in non-dominant hand precision drawing. Neuropsychologia, 87, 157–168. 10.1016/j.neuropsychologia.2016.05.016

Philip, B. A., Li, F., Hawkins-Chernof, E., Swamidass, V., & Zwir, I. (2023). Motor Assessment With the STEGA iPad App to Measure Handwriting in Children. American Journal of Occupational Therapy, 77(3).

Philip, B. A., Thompson, M. R., Baune, N. A., Hyde, M., & Mackinnon, S. E. (2022). Failure to Compensate: Patients With Nerve Injury Use Their Injured Dominant Hand, Even When Their Nondominant Is More Dexterous. Arch Phys Med Rehabil, 103(5), 899–907. 10.1016/j.apmr.2021.10.010

Picard, N., & Strick, P. L. (2001). Imaging the premotor areas. Current opinion in neurobiology, 11(6), 663–672.

Power, J. D., Mitra, A., Laumann, T. O., Snyder, A. Z., Schlaggar, B. L., & Petersen, S. E. (2014). Methods to detect, characterize, and remove motion artifact in resting state fMRI. NeuroImage, 84, 320–341.

Prichard, E., Propper, R. E., & Christman, S. D. (2013). Degree of Handedness, but not Direction, is a Systematic Predictor of Cognitive Performance. Front Psychol, 4, 9. 10.3389/fpsyg.2013.00009

Proske, U., & Gandevia, S. C. (2012). The proprioceptive senses: their roles in signaling body shape, body position and movement, and muscle force. Physiological Reviews.

Rolls, E. T., Deco, G., Huang, C.-C., & Feng, J. (2023). The human posterior parietal cortex: effective connectome, and its relation to function. Cerebral cortex, 33(6), 3142–3170.

Satterthwaite, T. D., Elliott, M. A., Gerraty, R. T., Ruparel, K., Loughead, J., Calkins, M. E., Eickhoff, S. B., Hakonarson, H., Gur, R. C., & Gur, R. E. (2013). An improved framework for confound regression and filtering for control of motion artifact in the preprocessing of resting-state functional connectivity data. NeuroImage, 64, 240–256.

Shen, J. (2022). Plasticity of the central nervous system involving peripheral nerve transfer. Neural plasticity, 2022(1), 5345269.

Smith, J. C., & Frey, S. H. (2011). Use of independent component analysis to define regions of interest for fMRI studies. International Society for Magnet Resonance in Medicine (ISMRM) Montreal.

Sokki, A. M., Bhat, D. I., & Devi, B. I. (2012). Cortical reorganization following neurotization: a diffusion tensor imaging and functional magnetic resonance imaging study. Neurosurgery, 70(5), 1305–1311.

Stone, K. D., & Gonzalez, C. L. (2015). Manual preferences for visually-and haptically-guided grasping. Acta Psychol (Amst), 160, 1–10. 10.1016/j.actpsy.2015.06.004

Taylor, K. S., Anastakis, D. J., & Davis, K. D. (2009). Cutting your nerve changes your brain. Brain, 132(11), 3122–3133.

Team, R. C. (2020). R: A language and environment for statistical computing. 2020. R Foundation for Statistical Computing: Vienna, Austria.

Tung, T. H., & Mackinnon, S. E. (2010). Nerve transfers: indications, techniques, and outcomes. The Journal of hand surgery, 35(2), 332–341.

Uswatte, G., Taub, E., Morris, D., Light, K., & Thompson, P. (2006). The Motor Activity Log-28: assessing daily use of the hemiparetic arm after stroke. Neurology, 67(7), 1189–1194.

Valyear, K. F., Mattos, D., Philip, B. A., Kaufman, C., & Frey, S. H. (2019). Grasping with a new hand: Improved performance and normalized grasp-selective brain responses despite persistent functional changes in primary motor cortex and low-level sensory and motor impairments. Neuroimage, 190, 275–288.

Vanderwal, T., Eilbott, J., & Castellanos, F. X. (2019). Movies in the magnet: Naturalistic paradigms in developmental functional neuroimaging. Dev Cogn Neurosci, 36, 100600. 10.1016/j.dcn.2018.10.004

Wandell, B. A., Dumoulin, S. O., & Brewer, A. A. (2007). Visual field maps in human cortex. Neuron, 56(2), 366–383.

Woo, C. W., Krishnan, A., & Wager, T. D. (2014). Cluster-extent based thresholding in fMRI analyses: pitfalls and recommendations. NeuroImage, 91, 412–419. 10.1016/j.neuroimage.2013.12.058

Worsley, K. J., Marrett, S., Neelin, P., Vandal, A. C., Friston, K. J., & Evans, A. C. (1996). A unified statistical approach for determining significant signals in images of cerebral activation. Human brain mapping, 4(1), 58–73.

Wu, J., Yan, T., Zhang, Z., Jin, F., & Guo, Q. (2012). Retinotopic mapping of the peripheral visual field to human visual cortex by functional magnetic resonance imaging. Human brain mapping, 33(7), 1727–1740.

Yang, Y. L. T.,; Deng, Y.; Wang, J.; Li, Y.; Liu, H.; Wang, W. (2024). Dynamic alternations of interhemispheric functional connectivity in brachial plexus avulsion injury patients with nerve transfer: a resting state fMRI study. Cereb Cortex, 34(1). 10.1093/cercor/bhad415

Yao, J., Chen, A., Kuiken, T., Carmona, C., & Dewald, J. (2015). Sensory cortical re-mapping following upper-limb amputation and subsequent targeted reinnervation: A case report. Neuroimage Clin, 8, 329–336. 10.1016/j.nicl.2015.01.010

Yu, A., Wang, S., Cheng, X., Liang, W., Bai, R., Xue, Y., & Li, W. (2017). Functional connectivity of motor cortical network in patients with brachial plexus avulsion injury after contralateral cervical nerve transfer: a resting-state fMRI study. Neuroradiology, 59(3), 247–253. 10.1007/s00234-017-1796-0

Zhao, W., Makowski, C., Hagler, D. J., Garavan, H. P., Thompson, W. K., Greene, D. J., Jernigan, T. L., & Dale, A. M. (2023). Task fMRI paradigms may capture more behaviorally relevant information than resting-state functional connectivity. NeuroImage, 270, 119946. 10.1016/j.neuroimage.2023.119946

