## Supplementary material for "Functional connectivity during drawing after upper extremity peripheral nerve surgery: enhanced connectivity between motor and visuomotor-parietal regions": Supplementray_Text.docx

**Supplementary Text**

**fMRI Preprocessing**

The data were preprocessed using fMRIPrep 24.1.1 (Esteban et al., 2019; RRID:SCR_016216), which is based on Nipype 1.8.6 (Gorgolewski et al., 2011; RRID:SCR_002502). Many internal operations of fMRIPrep use Nilearn 0.10.4 (Abraham et al., 2014; RRID:SCR_001362), mostly within the functional processing workflow.

**Anatomical preprocessing***:* The T1-weighted (T1w) volume was corrected for intensity non-uniformity with N4BiasFieldCorrection (Tustison et al., 2010), distributed with ANTs 2.5.3 (Avants et al., 2008; RRID:SCR_004757). The T1w-image was skull-stripped using antsBrainExtraction.sh workflow (using OASIS30ANTs template). The segmentation of cerebrospinal fluid (CSF), white-matter (WM) and gray-matter (GM) was performed on the brain-extracted T1w using fast (FSL, RRID:SCR_002823,(FSL, Zhang et al., 2002; RRID:SCR_002823). Brain surfaces were reconstructed using recon-all (FreeSurfer 7.3.2, Dale et al., 1999; RRID:SCR_001847), and the brain mask was refined by reconciling ANTs- and FreeSurfer-derived segmentations of the cortical gray-matter of Mindboggle (Klein et al., 2017; RRID:SCR_002438). Finally, the T1w volume was normalized to symmetrical MNI standard space (ICBM 152 Nonlinear Asymmetric template version 2009c [RRID:SCR_008796; TemplateFlow ID: MNI152NLin2009cAsym]) using nonlinear registration with antsRegistration (ANTs 2.5.3), producing an anatomical reference for functional co-registration.

**Functional preprocessing:** Each of the three BOLD functional runs was motion-corrected by estimating rigid-body transformations to a reference volume using mcflirt (FSL, Jenkinson et al., 2002). The estimated fieldmap was then aligned with rigid-registration to the target EPI (echo-planar imaging) reference run. The field coefficients were mapped on to the reference EPI using the transform. The BOLD reference was then co-registered to the T1w reference using bbregister (Freesurfer, Greve & Fischl, 2009). Several confounding time-series were calculated based on the preprocessed BOLD: framewise displacement (FD) along with the WM, CSF, and global signal. FD was computed using fsl_motion_outliers (relative root mean square displacement between affines, Jenkinson et al., 2002). The WM and CSF signals were extracted from masks derived from the brain segmentations, and the global signal from the whole brain mask. Three global signals were extracted from CSF, the WM, and the whole-brain masks. Additionally, a set of physiological regressors were extracted to allow for component-based noise correction (CompCor, Behzadi et al., 2007). Principal components were estimated after high-pass filtering the preprocessed BOLD time-series (using a discrete cosine filter with 128s cut-off) for the two CompCor variants: temporal (tCompCor) and anatomical (aCompCor). tCompCor components were then calculated from the top 2% variable voxels within the brain mask. For aCompCor, three probabilistic masks (CSF, WM and combined CSF+WM) were generated in anatomical space. These masks were resampled into BOLD space and binarized by thresholding at 0.99. The head-motion estimates and CSF, WM, and global signals were expanded with the inclusion of temporal derivatives and quadratic terms for each (Satterthwaite et al., 2013). Additional nuisance timeseries were calculated by means of principal components analysis of the signal found within a thin band (crown) of voxels around the edge of the brain, as proposed by Patriat et al. (2017). All resamplings were performed with a single interpolation step by composing all the pertinent transformations (i.e. head-motion transform matrices, susceptibility distortion correction when available, and co-registrations to anatomical and output spaces). Gridded (volumetric) resamplings were performed using nitransforms, configured with cubic B-spline interpolation.

Abraham, Pedregosa, Eickenberg, Gervais, Mueller, Kossaifi, Gramfort, Thirion, & Varoquaux. (2014). Machine learning for neuroimaging with scikit-learn. *Frontiers in neuroinformatics*, *8*, 14.

Avants, Epstein, Grossman, & Gee. (2008). Symmetric diffeomorphic image registration with cross-correlation: evaluating automated labeling of elderly and neurodegenerative brain. *Medical image analysis*, *12*(1), 26-41.

Behzadi, Restom, Liau, & Liu. (2007). A component based noise correction method (CompCor) for BOLD and perfusion based fMRI. *NeuroImage*, *37*(1), 90-101.

Dale, Fischl, & Sereno. (1999). Cortical surface-based analysis: I. Segmentation and surface reconstruction. *NeuroImage*, *9*(2), 179-194.

Esteban, Markiewicz, Blair, Moodie, Isik, Erramuzpe, Kent, Goncalves, DuPre, & Snyder. (2019). fMRIPrep: a robust preprocessing pipeline for functional MRI. *Nature methods*, *16*(1), 111-116.

Gorgolewski, Burns, Madison, Clark, Halchenko, Waskom, & Ghosh. (2011). Nipype: a flexible, lightweight and extensible neuroimaging data processing framework in python. *Frontiers in neuroinformatics*, *5*, 13.

Greve, & Fischl. (2009). Accurate and robust brain image alignment using boundary-based registration. [Evaluation Study]. *NeuroImage*, *48*(1), 63-72. <https://doi.org/10.1016/j.neuroimage.2009.06.060>

Jenkinson, Bannister, Brady, & Smith. (2002). Improved optimization for the robust and accurate linear registration and motion correction of brain images. *NeuroImage*, *17*(2), 825-841.

Karimpoor, Tam, Strother, Fischer, Schweizer, & Graham. (2015). A computerized tablet with visual feedback of hand position for functional magnetic resonance imaging. *Frontiers in Human Neuroscience*, *9*, 150.

Klein, Ghosh, Bao, Giard, Häme, Stavsky, Lee, Rossa, Reuter, & Chaibub Neto. (2017). Mindboggling morphometry of human brains. *PLoS Computational Biology*, *13*(2), e1005350.

Patriat, Reynolds, & Birn. (2017). An improved model of motion-related signal changes in fMRI. *NeuroImage*, *144*, 74-82.

Philip, & Frey. (2014). Compensatory Changes Accompanying Chronic Forced Use of the Nondominant Hand by Unilateral Amputees. *The Journal of Neuroscience: the Official Journal of the Society for Neuroscience*, *34*(10), 3622-3631. <https://doi.org/10.1523/JNEUROSCI.3770-13.2014>

Philip, Li, Hawkins-Chernof, Swamidass, & Zwir. (2023). Motor Assessment With the STEGA iPad App to Measure Handwriting in Children. *American Journal of Occupational Therapy*, *77*(3).

Satterthwaite, Elliott, Gerraty, Ruparel, Loughead, Calkins, Eickhoff, Hakonarson, Gur, & Gur. (2013). An improved framework for confound regression and filtering for control of motion artifact in the preprocessing of resting-state functional connectivity data. *NeuroImage*, *64*, 240-256.

Tam, Churchill, Strother, & Graham. (2011). A new tablet for writing and drawing during functional MRI. *Human Brain Mapping*, *32*(2), 240-248.

Tustison, Avants, Cook, Zheng, Egan, Yushkevich, & Gee. (2010). N4ITK: improved N3 bias correction. *IEEE transactions on medical imaging*, *29*(6), 1310-1320.

Zhang, Brady, & Smith. (2002). Segmentation of brain MR images through a hidden Markov random field model and the expectation-maximization algorithm. *IEEE transactions on medical imaging*, *20*(1), 45-57.
