## Supplementary material for "Functional connectivity during drawing after upper extremity peripheral nerve surgery: enhanced connectivity between motor and visuomotor-parietal regions": Table_S1.docx

| **Supplementary Table 1**. Nerve Injury Characteristics | | | | | |
| --- | --- | --- | --- | --- | --- |
|  | n (%) | | | Between Groups | |
| Variable | Repair  (n = 8) | Transfer  (n = 4) | Total  (n=12) | Statistic | *p* |
| Injured Nerve |  |  |  |  | 0.95 |
| Cutaneous | 1 (5.9) | 0 | 1 (4.3) |  |  |
| Median | 5 (29.4) | 1 (16.7) | 6 (26.1) |  |  |
| Posterior Interosseous | 1 (5.9) | 0 | 1 (4.3) |  |  |
| Radial | 2 (11.8) | 1 (16.7) | 3 (13) |  |  |
| Ulnar | 6 (35.3) | 3 (50) | 9 (39.1) |  |  |
| Other | 2 (11.8) | 1 (16.7) | 3 (13) |  |  |
| Injury Level |  |  |  |  | 1 |
| Branch/Thoracic Outlet | 1 (5.3) | 0 | 1 (3.6) |  |  |
| Elbow | 4 (21.1) | 2 (22.2) | 6 (21.4) |  |  |
| Forearm | 3 (15.8) | 2 (22.2) | 5 (17.9) |  |  |
| Wrist | 7 (36.8) | 3 (33.3) | 10 (35.7) |  |  |
| Hand | 3 (15.8) | 2 (22.2) | 5 (17.9) |  |  |
| Digit | 1 (5.3) | 0 | 1 (5.3) |  |  |
| Injury Cause |  |  |  |  | 0.23 |
| Injury | 5 (55.6) | 1 (20) | 6 (42.9) |  |  |
| Chronic Compression | 4 (44.4) | 2 (40) | 6 (42.9) |  |  |
| Surgical Complication | 0 | 2 (40) | 2 (14.3) |  |  |
| Tumor | 0 | 0 | 0 |  |  |
| Injury Severity |  |  |  |  | 0.01 |
| Neurapraxia | 6 (75) | 0 | 6 (50) |  |  |
| Axonotmesis | 0 | 3 (75) | 3 (50) |  |  |
| Neurotmesis | 2 (22.2) | 1 (25) | 3 (50) |  |  |
| # of OT prescriptions, *median* [range] | 1 [0-4] | 0.50 [0-2] |  | W=20.5 | 0.47 |
| # of PT prescriptions, *median* [range] | 0.50 [0-4] | 1.50 [1-2] |  | W=9 | 0.25 |
| Time since injury, *median* [range] | 61.50 [3-425] | 26 [10-158] |  | W=18 | 0.79 |
| Time since surgery, *median* [range] | 9 [1-418] | 16.50 [2-158] |  | W=14 | 0.79 |
| *Note*. Percentages may not equal 100% because individual patients could have multiple injured nerves, levels, and causes. Abbreviations: OT = occupational therapy; PT = physical therapy. Categorical variables were compared between groups using Fisher’s exact test due to cell counts <5. | | | | | |
