## Supplementary material for "Functional connectivity during drawing after upper extremity peripheral nerve surgery: enhanced connectivity between motor and visuomotor-parietal regions": Table_S2.docx

| **Supplementary Table 2**. Surgery Details | | | | | |
| --- | --- | --- | --- | --- | --- |
| Subject  ID | Group | # of Surgeries | Primary Procedure (most recent surgery) | Previous Procedures | Months Since Surgery |
| 1004 | Repair | 2 | Ulnar nerve repair (forearm)  Flexor tendon repair | Tenolysis | 418 |
| 1007 | Repair | 1 | Ulnar nerve transposition (elbow)  Flexor–pronator lengthening |  | 6 |
| 1011 | Repair | 1 | Ulnar nerve decompression (Guyon’s canal)  Ganglion decompression (Guyon's canal) |  | 2 |
| 1026 | Repair | 2 | Ulnar nerve repair (elbow)  Right shoulder rotator cuff | Median nerve release (carpal tunnel)  Ulnar nerve release (Guyon’s canal and cubital tunnel)  Decompression of nerve branches at elbow | 26 |
| 1032 | Repair | 1 | Posterior interosseous nerve decompression (forearm)  Radial sensory nerve decompression  Extensor tenotomies; de Quervain’s release |  | 12 |
| 1036 | Repair | 5 | Ulnar nerve transposition (elbow) | Thoracic outlet decompressions ×3  Ulnar nerve decompression (Guyon’s canal)  Carpal tunnel release; flexor–pronator lengthening | 23 |
| 1038 | Repair | 1 | Median and ulnar nerve decompression (wrist/Guyon’s)  Cubital tunnel release |  | 1 |
| 1051 | Repair | 1 | Median nerve repair (wrist, grafted) | Palmar cutaneous nerve graft; carpal tunnel release; FPL repair; FCR repair; skin graft | 3 |
| 1009 | Transfer | 2 | Anterior interosseous to ulnar motor nerve transfer | Cubital and carpal tunnel release; tendon tenodesis | 2 |
| 1023 | Transfer | 1 | Radial to median nerve transfer |  | 158 |
| 1027 | Transfer | 3 | Ulnar nerve transposition (elbow) | Sensory transfer to small finger  Tendon transfers for pinch/clawing | 30 |
| 1046 | Transfer | 2 | Anterior interosseous to ulnar motor nerve transfer | Ulnar nerve transposition; neurolysis; Guyon’s canal release | 3 |
| *Note.* # of Surgeries = Number of separate surgical events. Months Since Surgery = Months between fMRI scans and most recent surgery. | | | | | |
