## Supplementary material for "Functional connectivity during drawing after upper extremity peripheral nerve surgery: enhanced connectivity between motor and visuomotor-parietal regions": Table_S3.docx

| **Supplementary Table 3**. List of Activity Card Sort (ACS) Precision Hand Activities | |
| --- | --- |
| ACS Domain | Activity |
| Domestic Activity & Chores | Mending or repairing clothes |
| Domestic Activity & Chores | Fixing things around the house or grounds |
| Domestic Activity & Chores | Car maintenance |
| Domestic Activity & Chores | Paying bills |
| Domestic Activity & Chores | School work or assignments |
| Leisure Activity | Sewing (clothes or household items) |
| Leisure Activity | Quilting |
| Leisure Activity | Cards, including poker or solitaire |
| Leisure Activity | Board games |
| Leisure Activity | Needlecraft |
| Leisure Activity | Hand crafts |
| Leisure Activity | Jigsaw or other puzzles |
| Leisure Activity | Crossword puzzles or Sudoku |
| Leisure Activity | Arts (for example, painting, drawing, sculpture, weaving) |
| Leisure Activity | Playing a musical instrument |
| Leisure Activity | Creative writing, journal, or poetry |
| Leisure Activity | Letter writing (not including emails) |
| Leisure Activity | Taking photographs |
| Leisure Activity | Playing video games on a computer or console |
| Leisure Activity | Playing games on a smartphone |
| Active Leisure / Fitness | Hunting |
| Active Leisure / Fitness | Fishing |
| Active Leisure / Fitness | Rock Climbing |
| Social Interaction Activities | Text messaging family and friends |
| Social Interaction Activities | Board games or other group games (for example: chess, pool, Settlers of Catan, Taboo) |
| Social Interaction Activities | Email or instant messaging (aside from phone text messaging) |
