## Supplementary material for "Functional connectivity during drawing after upper extremity peripheral nerve surgery: enhanced connectivity between motor and visuomotor-parietal regions": Table_S4.docx

| **Supplementary Table 4.** Patients who provided subjective comments on Motor Activity Log | | | | | | | | | |
| --- | --- | --- | --- | --- | --- | --- | --- | --- | --- |
| **Motor Activity Log (MAL)** | | | | **Other characteristics** | | | | | |
| **Subject ID** | **Group** | **Amount of Use (0-5)** | **Participant Comment** | **Months Since Injury** | **Months Since Surgery** | **ΔEHI** | **Ability** | **Retention (%)** | **RH Use** |
| 1009 | Transfer | **5.0** | Reports difficulty with tasks and functional loss affecting golf grip and removing small items from the pocket, not fully reflected in questionnaire | 10 | 2 | 0 | 89.16 | 100 | 0.75 |
| 1011 | Repair | **4.65** | Patient wanted to note that they consider the actions more intentional rather than automatic "hesitates" | 5 | 2 | 0 | 92.24 | 58.33 | 0.61 |
| 1023 | Transfer | **4.5** | Has been living with the condition so long and compensates by using his left hand | 158 | 158 | 0 | 77.5 | 72.72 | 0.40 |
| 1026 | Repair | **4.06** | Bimanual stuff is difficult like hair-dryer with round brush | 50 | 26 | -70 | 48.33 | 84.62 | 0.41 |
| 1036 | Repair | 2.42 | Uses adapted pen for writing | 112 | 23 | -115 | 67.5 | 66.66 | 0.17 |
| 1051 | Repair | 2.28 | Generally, forces themself to complete tasks as part of continued daily therapy. No tactile feedback, replaces physical "feel" with watching the hand do the activity | 3 | 3 | -75 | 79.16 | 81.25 | 0.05 |
| *Note.* Shaded = Participants with discordance between MAL amount of use and MAL optional subjective comments. “Other characteristics” are reproduced from Table 2 for side-by-side comparison with participant comments. | | | | | | | | | |
