## Supplementary figures and images for "Functional connectivity during drawing after upper extremity peripheral nerve surgery: enhanced connectivity between motor and visuomotor-parietal regions"

### SuppFig1.png

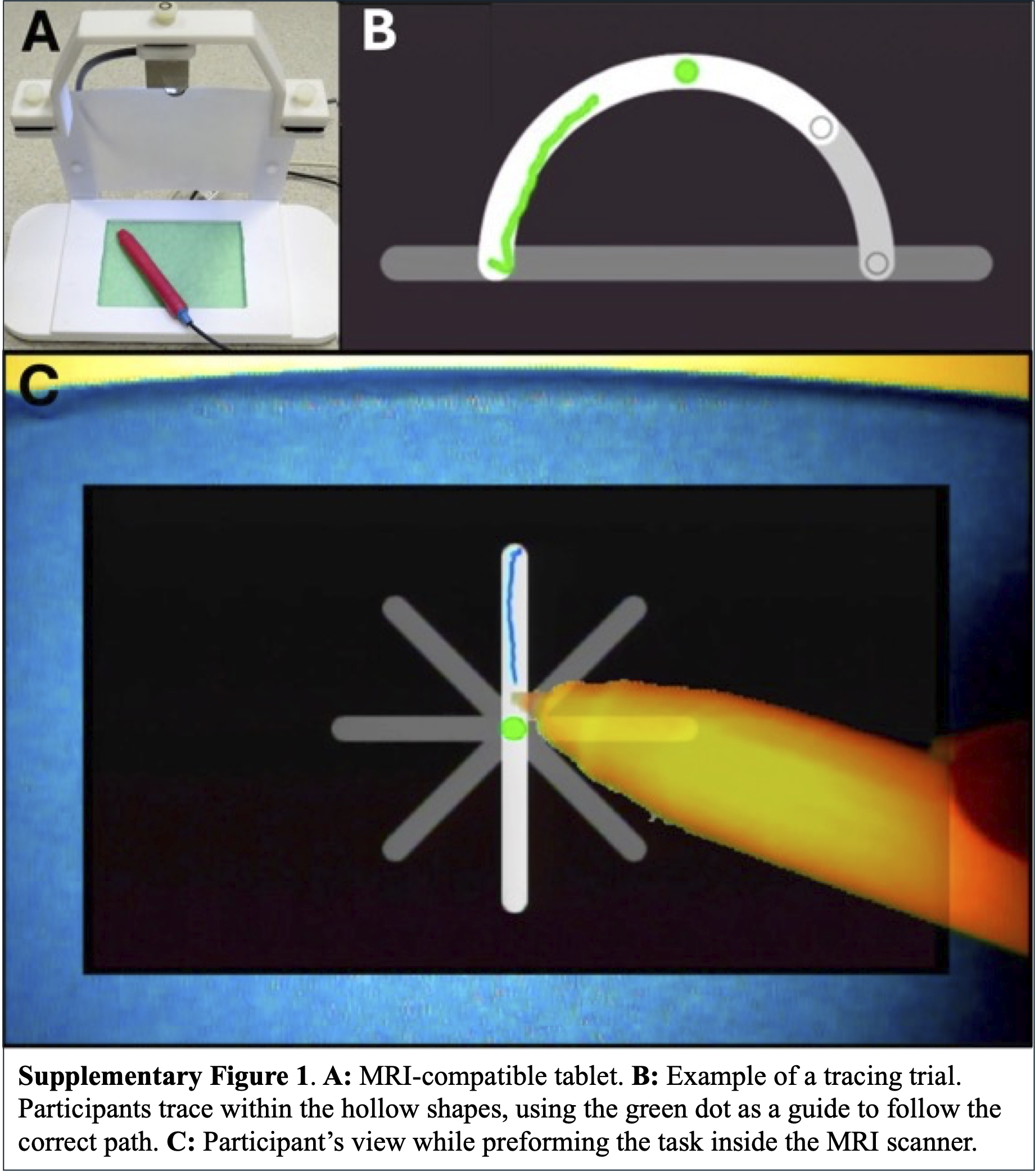

### SuppFig2.png

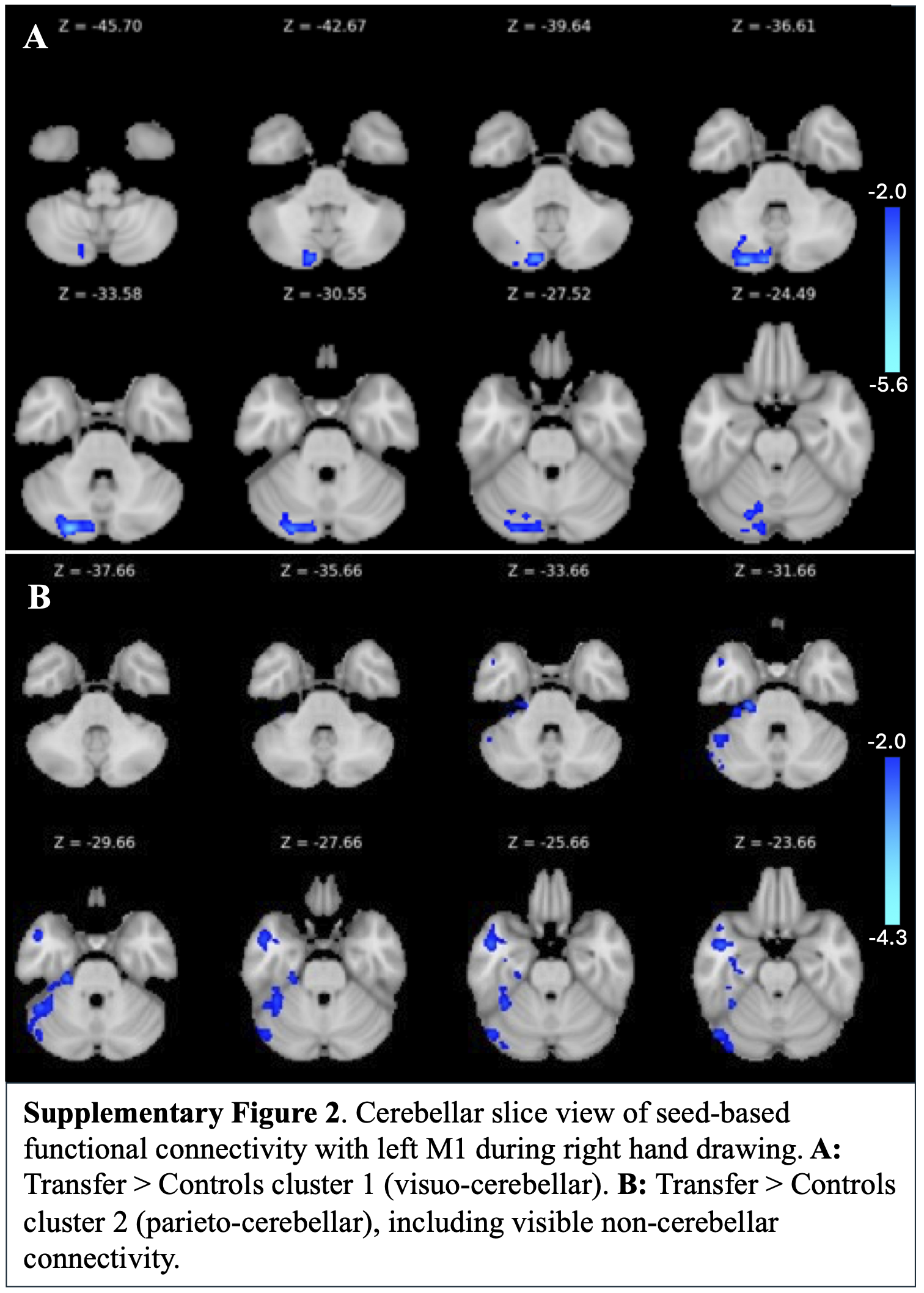
